# Synthetic communities as a model for determining interactions between a biofertilizer chassis organism and native microbial consortia

**DOI:** 10.1101/2025.02.13.638169

**Authors:** Cody S. Madsen, Jeffrey A. Kimbrel, Patrick Diep, Dante P. Ricci

## Abstract

Biofertilizers are critical for sustainable agriculture since they can replace ecologically disruptive chemical fertilizers while improving the trajectory of soil and plant health. Yet, to continue improving deployment, the persistence of designer biofertilizers within native soil consortia must be elucidated and enhanced. Here, we describe a high-throughput, modular, and automation-friendly *in vitro* approach to screen for biofertilizer organism persistence within soil-derived consortia after co-cultivation with stable synthetic soil microbial communities (SynComs) obtained through a top-down cultivation process. We profiled ∼1200 SynComs isolated from various soil sources and cultivated in divergent media types, and detected significant phylogenetic diversity (e.g., Shannon index > 4) and richness (Observed richness > 400) across these communities. We observed high reproducibility in SynCom community structure from common soil and media types, which provided a testbed for assessing biofertilizer persistence within representative native consortia. Furthermore, we demonstrate the screening method described herein can be coupled with microbial engineering to efficiently identify soil-derived SynComs where an engineered biofertilizer organism (i.e. *Bacillus subtilis*) persists. Additionally, our approach enables an analysis of the ecological impact of *B. subtilis* inoculation on SynCom structure and profile alterations in community diversity and richness (or lack thereof) associated with the presence of a genetically modified model bacterium. Ultimately, this work establishes a modular pipeline that could be integrated into a variety of microbiology/microbiome-relevant workflows or related applications that would benefit from assessing persistence and interaction of a specific organism of interest with native consortia.

## Introduction

Biofertilizers are critical agents for sustainability in agriculture that offer several advantages over traditional fertilizers, including improved cost-effectiveness, natural promotion of long-term plant and soil health, and a more favorable environmental and ecological impact[1–6]. However, issues such as poor persistence in target soil, deleterious interactions with native microbiota, or low functional robustness can limit the effectiveness (and, therefore, adoption) of biofertilizers relative to chemical agents[2,5,6]. These liabilities could potentially be mitigated through the discovery or design of specialized biofertilizer microbes with optimal commensal properties, but the establishment and maintenance of a stable rhizosphere that includes such strains is likely to require information about how effector strains interact with target soil biomes. Although effective and generalizable methods exist for introducing biofertilizer microbes into a target soil (e.g. through seed coating or in-furrow application[5,7–11]), detailed information about microbiome dynamics following biofertilizer introduction in native soils is largely limited to highly specific conditions[9] or is privately held by for-profit entities[12]. Understanding how field-deployable custom biofertilizers impact and interact with native microbiomes will be indispensable for the ultimate improvement of biofertilizer performance.

Recent investigations into microbial community dynamics have demonstrated the remarkable taxonomic diversity that can be successfully cultivated *in vitro* in the form of stable, reduced-complexity consortia using simple and cost-effective methods[13–15]. Further efforts have explored the impact of ecological engineering on the composition and diversity of *in vitro* communities and have illustrated experimental approaches to generating divergent communities from common source material[16,17]. Accordingly, these synthetic communities (SynComs) have become a potential new predictive tool in understanding biofertilizer impact and synergy with native consortia, although there is limited precedent for broad, longitudinal characterization of broadly diverse SynComs in natural ecosystems[18–20]. Additionally, SynComs can be isolated through holistic (top-down) and reductionist (bottom-up) approaches and have provided insights into rhizosphere dynamics, plant-microbe interactions, and invasion resistance, but the ecological principles that underlie assembly of durable communities and are still too poorly understood to enable effective *a priori* SynCom design[18,20–22]; this reflects, in part, the increasing need for novel methods that effectively and efficiently provide information on microbial community composition and activity with temporal resolution.

A particularly compelling approach to incorporating functional “effector” strains into divergent ecologies involves the development of synthetic biofertilizer microbial consortia containing microbes from the native target niche(s) that support stable maintenance of strains with desired functionality, e.g. through effective nutrient cycling or production of secondary metabolites[23–26]. One well-characterized candidate chassis for functional engineering and deployment is *Bacillus subtilis*, a Gram-positive facultative anaerobe and plant growth promoting rhizobacterium (PGPR) that has been shown to protect plants from environmental stress (e.g., ethylene toxicity, salt stress, toxic biotic agents[27,28]), form biofilms around plant roots for extended colonization, interact symbiotically with other microorganisms in the rhizosphere, resist extreme environmental conditions (e.g., heat, salt, extremes of pH) with and without spore formation[29–31] and enhance growth of cereals and legumes even in marginal soils[31–33]. *B. subtilis* is generally recognized as safe (GRAS) [34–39] and is used routinely in synthetic biology applications; importantly, genetic tools responsive to rhizosphere-relevant signals (sugars such as xylose [40–42] and mannose, elevated salt concentrations, or thermal energy) have been successfully developed for and characterized in *B. subtilis*[43–47]. In addition to PGPR function, *B. subtilis* is used for industrial production with demonstrated scalability and commercialization potential[35–37]. *B. subtilis* could therefore represent an ideal chassis organism for future biofertilizer technology and a tractable tool for characterizing the interactions of engineered organisms with native soil consortia.

SynComs are predicted to be a valuable tool in understanding microbial ecology and enabling sustainable agriculture, but best practices for developing and deploying such systems are yet to be defined[48–50]. Therefore, we aimed to build on recent top-down approaches to SynCom cultivation[51] and establish new modular pipelines that could be used to screen for desired functions and assess biofertilizer organism persistence with native consortia. We developed a high-throughput workflow to screen for the durable persistence of a microbe of interest following introduction into diverse soil-derived communities (Fig. 1, Supp. Fig. 1-2). We show that the presence of *B. subtilis* in highly divergent complex consortia can be readily detected through straightforward phenotypic assays amenable to high throughput screening and demonstrate persistence of *B. subtilis* into native soil derived consortia. We also uncover the impact of *B. subtilis* on community structure and propose that the pipeline reported here represents a useful platform for characterizing biofertilizer organism interaction with native consortia.

**Figure 1.**
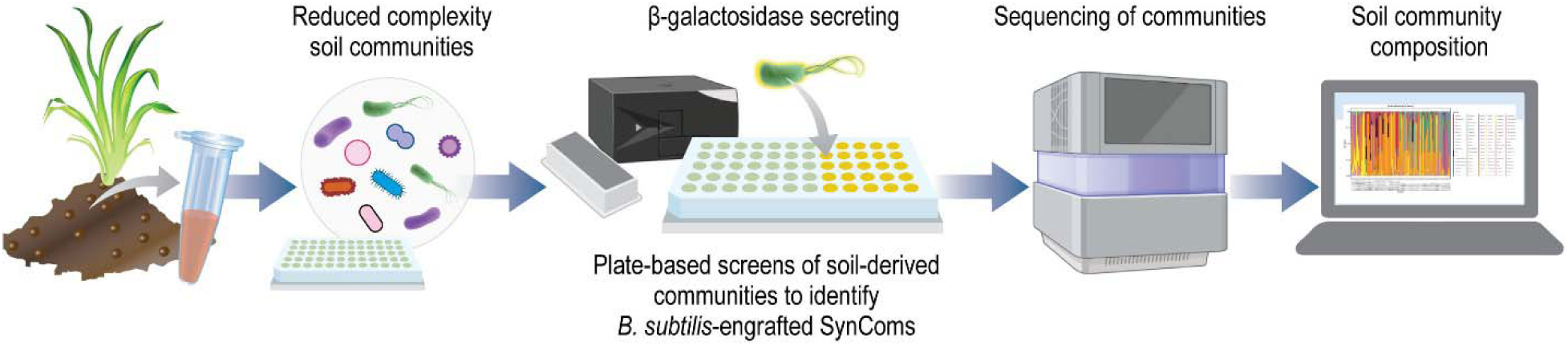
A high-throughput workflow for developing top-down SynComs and testing engineered microbe persistence. Reduced complexity SynComs were generated from multiple soils and various media types to represent native consortia then these SynComs were used as a platform for testing engineered *B. subtilis* ability to interact and persist with the native consortia.

## Results

### Top-down SynComs were created to represent native consortia

A top-down approach represents a straightforward and mechanism-agnostic way (i.e., microbes that sustainably interact will naturally propagate) to derive communities that represent some of the complexity in native ecology. Therefore, in the interest of developing a broad array of candidate soil-derived communities for testing persistence of engineered *B. subtilis*, we employed a top-down approach to cultivate microbial communities derived from three distinct soil sources in six different media conditions each. Furthermore, we reasoned that propagating communities from separate aliquots of the same soil used to make replicate 96-well plates would aid in discerning reproducibility from each soil type and test deterministic principles, so this was repeated for all three soil types. The media types included minimal (Cold Spring Harbor standard M9) and rich media (nutrient broth—NB, tryptic soy broth—TSB) to propagate a diverse portfolio of communities. Succinate was paired with glucose in half of the conditions to test how the two primary carbon sources from opposite ends of the metabolic cycle (EMP pathway and TCA cycle) could help construct sustained interactions between diverse community members[51]. Additionally, succinate is one of the primary carbon sources utilized by microbes when interacting with the plant host so this could increase cultivation of microbes that biofertilizers would need to persist with in the soil[52,53]. Back-dilution was repeated ten times[51] to create stability in the communities which were then carried through two freeze thaw cycles then back-diluted three more times before proceeding to preparation for 16S rRNA amplicon sequencing (Supp. Fig. 1).

The top-down approach was determined to produce diverse and rich SynComs that varied based on soil, media type and 96-well plate replicate after 400 communities were sequenced (Fig. 2). When assessed at the soil level, Hopland soil[54,55] (northern California pasture soil) and Potting soil (Home Depot; Kellogg Garden Organics Indoor Potting Mix) had greater diversity and richness compared to soil sampled from Lawrence Livermore National Laboratory (LLNL) based on several alpha diversity and richness metrics and were comparable to or greater than the community standard (Supp. Fig. 3). The beta diversity metrics showed the communities clustered based on soil type when visualized using principal component analysis (PCA) plots, and the same result was observed when a phylogenetic tree at the community level was constructed (Supp. Fig. 3, 4a). Overall, the communities from distinct soil sources yielded compositionally distinct communities even when propagated in the same media conditions. In contrast, SynComs grown in different media types produced further unique communities of varying diversity and composition. Specifically, the richer media and media that contained succinate generally exhibited greater diversity and richness, but this varied according to the original soil source (Fig. 2, Supp. Fig. 5, 6A). Based on the beta diversity metrics, phylogenetic tree and composition profile, the media types were less clustered and more generally distributed than the more distinct clustering and impact from the original soil source (Fig. 2, Supp. Fig. 4B, 5, 6B). Yet, even with variance observed within and across soil types, consistency in the form of similar composition and relative abundance was observed within replicates grown independently in the same media type from the same soil source (i.e. technical replicates; Supp. Fig. 6B).

**Figure 2.**
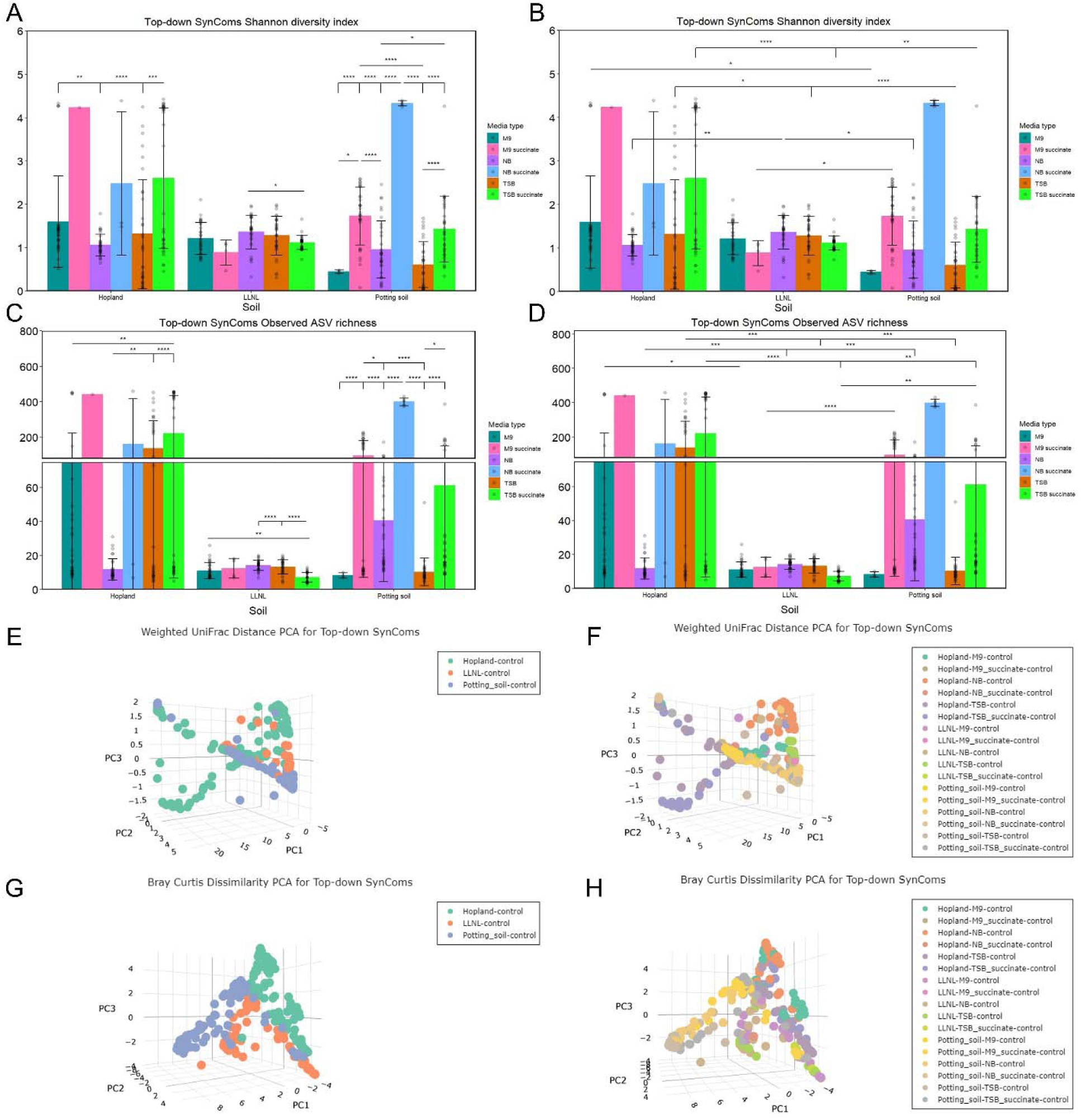
Diverse and rich top-down SynComs were propagated with differences and grouping related to soil and media type. After 16S rRNA sequencing base or control top-down SynComs, several metrics were calculated to assess the SynComs. Shannon diversity index (A, B) and Observed amplicon sequencing variant (ASV) richness (C, D) are shown for estimating alpha diversity and richness respectively with intra-soil statistics shown on the left (A, C) and inter-soil statistics shown on the right (B, D). For beta diversity, Weighted UniFrac distance (E, F) and Bray Curtis Dissimilarity (G, H) are visualized as PCA plots with coloring by soil on the left (E, G) and coloring by soil and media type on the right (F, H). The designation of control is applied to these SynComs as these are the base SynComs that the two *B. subtilis* strains were inoculated into generating the other SynComs. Data for alpha diversity is mean ± SD with each SynCom represented as a point which is up to 32 individual replicates as there were 16 SynComs in each media type of every soil and two separate 96-well plates were sequenced for every condition with the same for PCA plots; *p<0.05, **p<0.01, ***p<0.001, ****p<0.0001.

However, when comparing different 96-well plates stemming from sub-samplings of the same soil source (i.e., biological replicates), divergence in communities even in the same media type was demonstrated especially from the more diverse soil sources (Supp. Fig. 6C-7).

Diversity metrics quantified for various SynComs from Hopland and Potting soil showed their communities reached high levels of diversity based on Shannon scores being greater than 3 or 4, standardized effect size (SES) of mean pairwise distance (MPD) phylogenetic diversity z-values approaching 4, and Simpson scores nearing 1 (Fig. 2, Supp. Fig. 5, 7). Richness metric for the same communities paralleled the alpha diversity metrics with Observed amplicon sequencing variant (ASV) richness reaching into the hundreds consistently and observed with some SynComs nearly at 500 (Fig. 2, Supp. Fig. 5, 7). Composition profiles provided clarity on these metrics by showing that 48 unique families could be cultivated in just one well-plate from Hopland soil with some SynComs reaching over 10 unique families (Supp. Fig. 6B). Some primary families in Hopland soil included *Pseudomonadaceae, Yersiniaceae, Xanthomonadaceae, Lachnospiraceae* and *Lactobacillaceae* to name a few (Supp. Fig. 4b).

However, LLNL and Potting soil SynComs were dominated by the *Enterobacteriaceae* and *Pseudomonadaceae* families and Potting soil SynComs also mirrored Hopland family composition in some SynComs (Supp. Fig. 6A,C). Accordingly, the phylogenetic trees and composition profiles revealed the broad diversity, divergence and variance in community structure that was generated from a top-down SynComs approach (Supp. Fig. 4,6). Additionally, we compared a representative set of SynComs across the storage and regrowth process and determined that overall, the communities exhibited retention of diversity with some fluctuations in individual SynCom structure (Supp. Fig. 8). Three-dimensional visuals of beta diversity and selective composition profile visuals were generated to provide further context and comparisons beyond the two-dimensional visuals (GitHub, Code Availability statement).

### *B. subtilis* was engineered to be trackable during interaction with SynComs

In pursuit of our goal of developing synthetic soil-derived communities that allow persistence of a non-native engineered organism and lacking a reliable means to predict the composition of potentially compatible communities, we sought to perform a screen for *in vitro* SynComs in which an engineered derivative of the model organism *B. subtilis* can stably persist. We reasoned this could be achieved by inoculating all communities with engineered *B. subtilis* and directly screening the resulting consortia for a unique engineered phenotype specific to the added strain. To develop a phenotypic screen, we modified each of two common *B. subtilis* strains (168 and 6051a) to include a xylose-inducible[56–58] *lacZ* gene into the host chromosome, yielding strains that produce beta-galactosidase (β-gal) in the presence of xylose, and can be distinguished based on o-nitrophenyl-β-d-galactopyranoside (ONPG) colorimetric detection of β-gal activity with minimal impact on community dynamics. Importantly the engineered strains were designed to secrete[59] β-gal into the extracellular milieu, which enabled direct screening of supernatants without requiring lysis or cell permeabilization, significantly simplifying the screening methodology and in theory reducing background associated with endogenous enzyme activity in the soil-derived isolates.

Expression of β-gal from the *B. subtilis* strains was tested without SynComs but under the same parameters as would be used for screening when the strains were added to the SynComs to create associations between amount of *B. subtilis* and ONPG readout. Serial dilutions without SynComs were performed to create the calibration curves between the amount of *B. subtilis* present and the change in absorbance from β-gal activity in cleaving ONPG which also established contextualization for background absorbance from the baseline SynComs (i.e., when the absorbance change could be estimated to come from *B. subtilis* strains or from native consortia). Even under these poor expression conditions in the static 96-well assay plates, control of β-gal expression was observed in both strains using the native *B. subtilis* xylose inducible system and the 6051a strain caused a greater change in absorbance in lysogeny broth (Supp. Fig. 9). Accordingly, the serial dilution expression assays were repeated in all the media types used to generate the SynComs to create calibration curves for each media type (Fig. 3).

**Figure 3.**
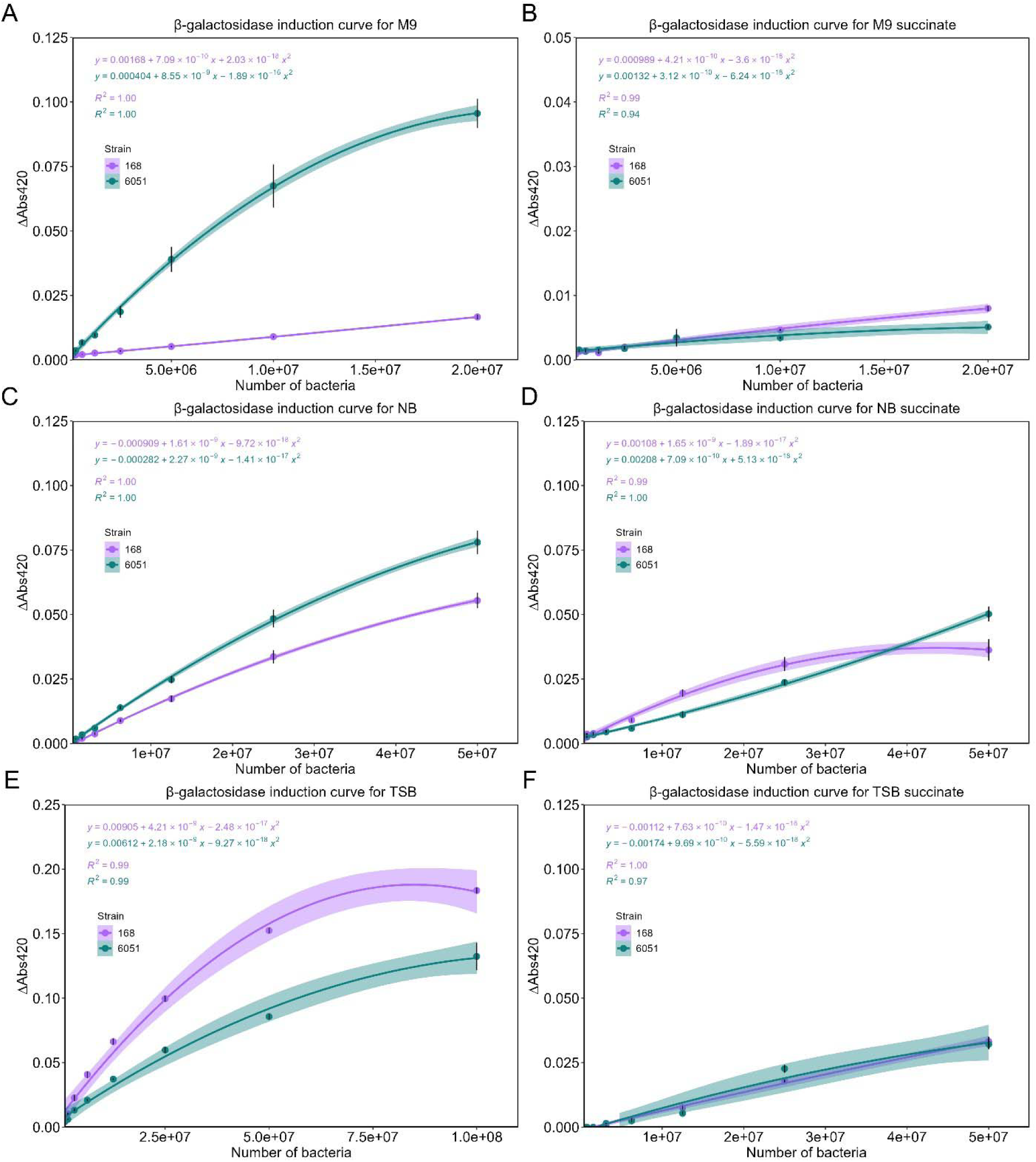
Calibration curves were created to connect *B. subtilis* strains abundance to change in absorbance from β-gal expression. *B. subtilis* 168 and 6051a (6051) strains expressing β-galactosidase under control of the xylose inducible system were serially diluted and induced with xylose in the conditions that would be used for screening in the SynComs to test change in absorbance after adding ONPG in all media types associated with SynCom growth. Best fit polynomial equations were generated in R to connect change in absorbance to abundance of each *B. subtilis* strain. Data is mean ± SEM of 12 replicates (n = 3 biological, n = 4 technical).

In all the media types, expression correlated with abundance of *B. subtilis* for both strains and 6051a generally caused a greater change in absorbance (Fig. 3). Additionally, in the media conditions containing succinate, the expression was lessened which indicated worse expression conditions and this was reinforced when the strains were grown in all the media types (Fig. 3, Supp. Fig. 10). Both strains’ growth rate was attenuated in the media conditions with succinate but both strains exhibited some growth in all conditions (Supp. Fig. 10).

### *B. subtilis* was tracked and identified in diverse communities from multiple soil types

To screen and downselect for SynComs that could contain the *B. subtilis* strains, the methods utilized to develop the calibration curves were translated into the community format. After the base or control top-down SynComs were propagated to stability, the SynComs were reanimated (i.e., grown up from glycerol stock after initial ten passages to establish SynComs) so the *B. subtilis* strains could be inoculated then tracked using ONPG. The *B. subtilis* strains were grown and normalized in optical density (OD_600_ = 1) then 5 µl was inoculated into the SynComs following the same passaging protocol as the base SynComs. Accordingly, the SynComs were outgrown and passaged as when the SynComs were created with screens performed at every passage to create a temporal tracking of changes in absorbance. Furthermore, the change in absorbance from the control top-down SynComs was subtracted from the change in absorbance in the inoculated SynComs for tracking ONPG change over time to screen for *B. subtilis* presence. Control top-down SynComs that produced absorbance values greater or less than the maximum value for each *B. subtilis* strain in each media type in the calibration curves were determined to be high or low background respectively. Accordingly, this process created categories to classify the SynComs after the ONPG screen which was used to downselect SynComs (same ones for both 168 and 6051a) with matching control SynComs for 16S rRNA sequencing. After sequencing, the four categories produced were persistence or no persistence with either high or low background with persistence being determined as consistent absorbance above background by *B. subtilis* inoculated SynComs combined with sequencing determining *Bacillus* ASV presence only in inoculated SynComs without any *Bacillus* ASV in the associated control SynComs. Lastly, the calibration curves were used to estimate a relative abundance tracking for *B. subtilis* in SynComs that were revealed to have persisted based on sequencing.

The ONPG screen proved successful to assess and downselect which SynComs may contain the *B. subtilis* strains (*viz.* SynComs which created absorbance changes consistently above background) especially in conditions where the control SynComs produced low levels of background. Ultimately, 400 SynComs from both 168 and 6051a inoculation conditions that matched the control top-down SynComs were selected and sequenced. After cross validating with sequencing, the ONPG screen identified key conditions where persistence occurred, as emphasized by the funnel charts depicting how the ONPG screen and sequencing categorized the SynComs (Fig. 4, Supp. Fig. 11-18). Specifically, for the 168 strain, the ONPG screen isolated the LLNL soil and the media types where the most persistence was seen for this strain (Fig. 4, Supp. Fig. 11A). As the conditions were subdivided within the LLNL soil, the ONPG temporal tracking curves for the well-plate and NB media type revealed significant separation in absorbance between the SynComs indicated to contain *B. subtilis* and those without *B. subtilis* (Fig. 4). Yet, other conditions such as other media types in LLNL or in Potting soil and Hopland soils that only contained one or a few SynComs with *B. subtilis* showed some separation, but the few representatives limited a trend from forming (Fig. 4G-H, Supp. Fig. 13, 15A-D, 16, 17A- D, 18A-B). Specifically, for the 6051a strain, a few trends of the SynComs containing *B. subtilis* separating from background were observed in both LLNL and Hopland in the specific media types where the majority of the 6051a strain persistence was present (Supp. Fig. 11, 14, 17E-H, 18C-D). Yet, this trend was not readily apparent in the Potting soil conditions for 6051a (Supp. Fig. 15E-H). Overall, the temporal tracking curves assisted in selecting for and identifying *B. subtilis* persistence with SynComs and highlighted a specific soil and media condition (Potting soil NB succinate) where both the 168 and 6051a strains persisted with very similar community structures (Supp. Fig. 16). The calibration curves were also used to convert the change in absorbance above background to relative abundance of the strains. The LLNL NB condition for the 168 strain demonstrated the utility of this process as the SynComs with the top relative abundance in the abundance tracking curves (NB 3, 4, 5, 9, 11) also had parallel results in sequencing based on the composition profile (Supp. Fig. 12A, C). These SynComs were associated with the low background conditions (Fig. 4) which overall connected to improved ability of the tracking curves to identify and quantify persistence in SynComs compared to other SynComs under low background conditions. Accordingly, *B. subtilis* containing SynComs in high background conditions or with very low abundance were difficult to quantify with the abundance curves (Supp. Fig. 12). Nonetheless, the temporal tracking and abundance curves did provide insight into the metabolic activity and stability of the *B. subtilis* strains in the SynComs over time instead of just an endpoint as with the sequencing. Ultimately, the *B. subtilis* strains persisted in diverse SynComs from all soil and media types with a variety of community structures as shown by phylogenetic trees, though the *B. subtilis* strains consistently persisted with specific genera such as *Pseudomonas* and *Klebsiella* of the native consortia across multiple soil and media types while not dominating community dynamics in any condition (Supp. Fig. 11).

**Figure 4.**
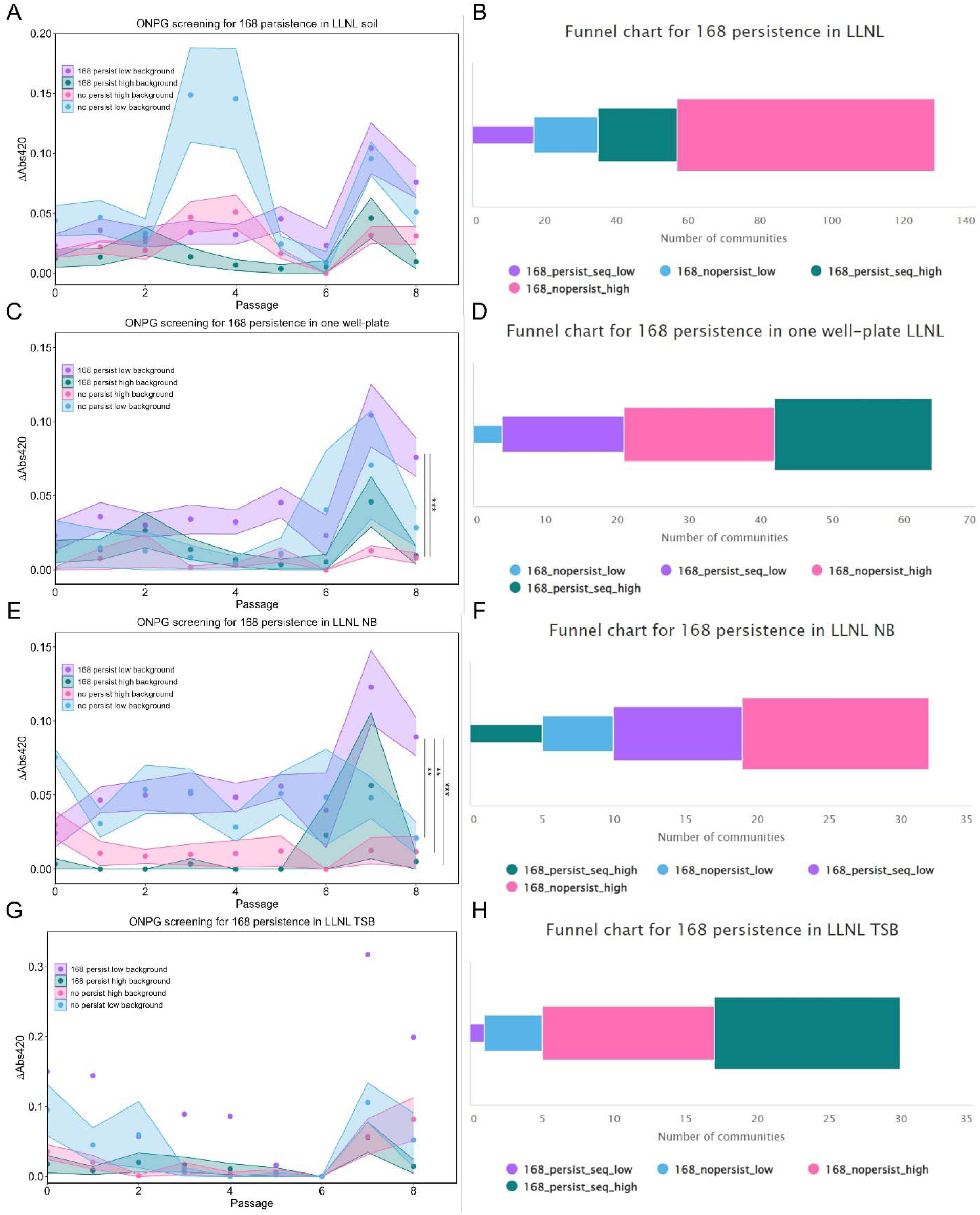
ONPG screen identified SynComs in specific soil and media conditions where the 168 strain expression of β-gal was distinguishable from background as a downselection mechanism for sequencing. Temporal tracking curves were constructed by plotting change in absorbance of SynComs that the 168 strain was inoculated into after subtracting background from the original base SynComs without inoculation. The SynComs were classified into four groups after persistence was indicated or not indicated by sequencing (persist or no persist). High or low background was determined by whether the original base SynCom was quantified to have absorbance values greater or less than the maximum value for each strain in each media type in the calibration curves. The tracking curves are plotted based on soil (A), then subdivided to just one well plate (C) or subdivided into two media types (E, G). Funnel charts (B, D, F, H) which are paired with the adjoining tracking curves are shown to represent the number of SynComs that were grouped into each subcategory based on the downselection process of the ONPG screen and sequencing. Colors for categories are the same between both types of plots (sequencing indicated—persist_seq, not indicated—nopersist; high background—_high, low background—_low). Points in the tracking curves is mean ± SEM of the number of replicates shown in the funnel charts based on how the SynComs were grouped; **p<0.01, ***p<0.001.

### SynComs were both altered and unaffected by inoculating *B. subtilis* process

After persistence was tracked and revealed by 16S rRNA sequencing, all the control SynComs and SynComs inoculated with the *B. subtilis* strains were further assessed using the metrics utilized for just the control SynComs. These metrics elucidated the impact on the SynComs from the inoculation process as all communities were sequenced simultaneously. Accordingly, the visualizations and statistics were structured to compare the control or base SynComs, 168 and 6051a inoculated SynComs to reveal any trends or specific outcomes.

By assessing all the SynComs’ alpha and beta diversity, composition profiles and phylogenetic trees, we quantified that the inoculation process altered the structure of SynComs in certain conditions, however some SynComs were unaffected by the inoculation process. In Potting soil and Hopland, the inoculation process significantly reduced diversity and richness in most of the media conditions except where the richness was reduced but the diversity did not significantly change (e.g., Potting soil TSB succinate, Hopland NB) or there was minimal change in diversity and richness as in Potting soil M9 (Fig. 5, Supp. Fig. 19-20A, B, I, 21). The beta diversity plots mirrored the alpha diversity and richness results as the soil and media conditions where the most significant changes were seen for alpha diversity and richness are the conditions where the PCA plots showed separation of the inoculation conditions compared to control such as Hopland TSB, Hopland TSB succinate and Potting soil NB succinate (Fig. 5, Supp. Fig. 20, 22). Yet, variance was observed in many Hopland and Potting soil conditions in multiple metrics and representative composition profiles confirmed that the inoculation process resulted in varying impact on the community structure between well-plates (Fig. 5, Supp. Fig. 19-20, 23). More broadly, in Hopland and Potting soil, the two strains had similar impact on SynCom structure in many conditions with some exceptions including Hopland TSB and TSB succinate where the 6051a strain was shown to persist in low abundance which denoted that persistence can be elucidated even from broad metrics (Fig. 5, Supp. Fig. 12, 20). Three-dimensional visuals of beta diversity and selective composition profile visuals were generated to provide further context and comparisons beyond the two-dimensional visuals (GitHub, Code Availability statement).

**Figure 5.**
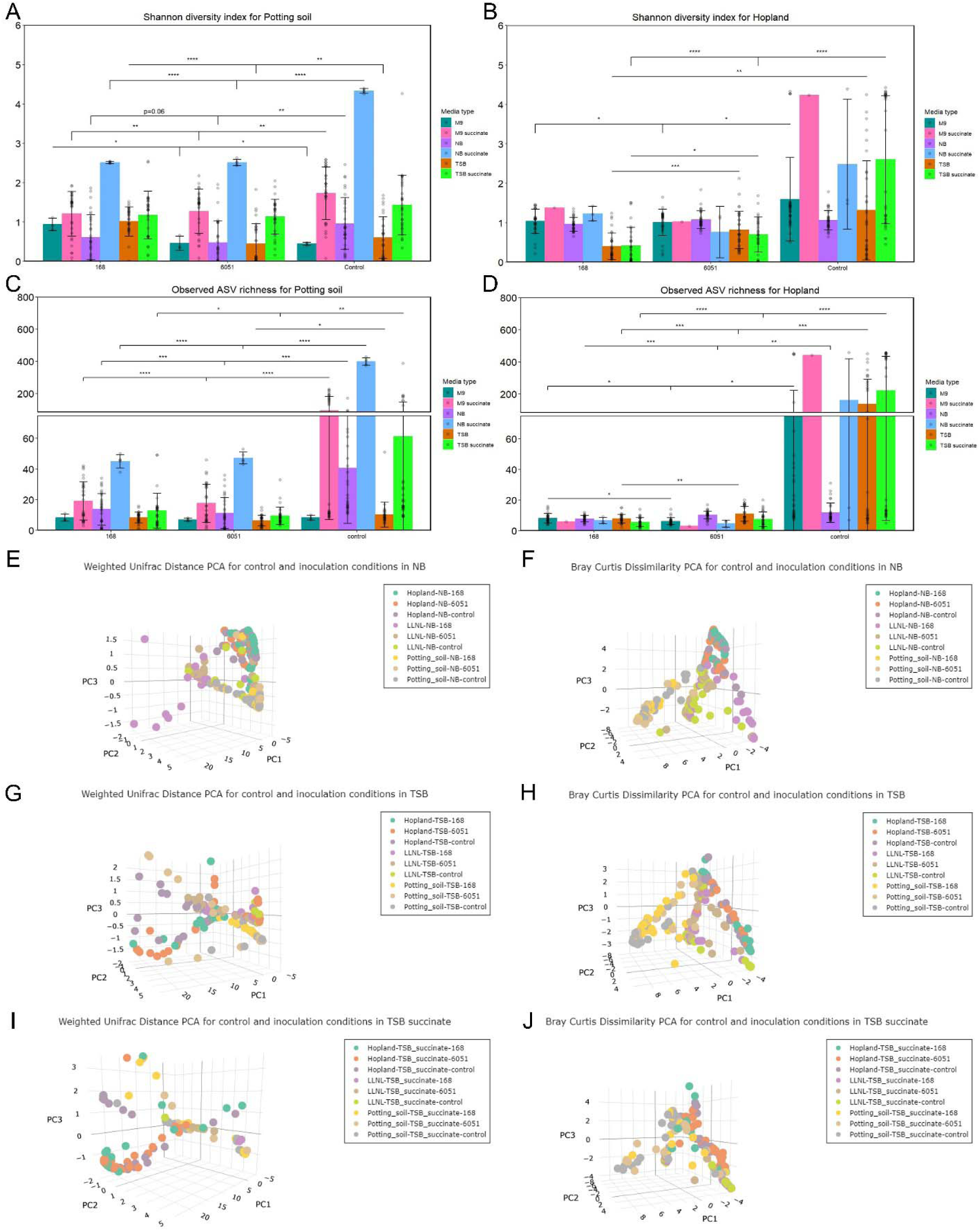
SynComs in Potting soil and Hopland were altered by inoculating *B. subtilis* 168 and 6051a strains. After analyzing all the SynComs (control or base and inoculation conditions) post 16S rRNA sequencing, several metrics were calculated to assess the change in the SynComs after inoculating the SynComs with 168 and 6051a (6051) strains. Shannon diversity index (A, B) and Observed amplicon sequencing variant (ASV) richness (C, D) are shown for alpha diversity and richness respectively with inter-inoculation statistics shown. For beta diversity, Weighted UniFrac distance (E, G, I) and Bray Curtis Dissimilarity (F, H, J) are visualized as PCA plots with graphs grouped based on media type then comparing soil and inoculation type. Data for alpha diversity is mean ± SD with each SynCom represented as a point which is up to 32 individual replicates as there were 16 SynComs in each media type of every soil and two separate 96-well plates were sequenced for every condition with the same for PCA plots; *p<0.05, **p<0.01, ***p<0.001, ****p<0.0001.

### *B. subtilis* appeared to fulfill environmental niche in SynComs from LLNL soil

While the Hopland and Potting soil SynComs inoculated with the *B. subtilis* strains exhibited many of the potential outcomes mentioned above (Fig. 5), minimal persistence and a reduction of SynCom complexity were the primary outcomes observed. However, for LLNL soil, the inoculation process had the opposite effect compared to Hopland and Potting soil as the process either increased or minimally impacted the diversity and richness of the SynComs (Fig. 6, Supp. Fig. 24). The beta diversity PCA plots revealed distinct separation of some SynComs with the clearest example being in the 168-peristence condition in LLNL NB (Fig. 6). These resulting trends towards increase in diversity, richness, and distance on the beta diversity PCAs were cross validated by composition profiles and these conditions correlated with the ONPG tracking and abundance curves as the most prominent persistence conditions (Fig. 6). The LLNL NB condition for the 168-persistence epitomized this result as mentioned above and this also altered the SynCom structure specifically in the 168-persistence (Fig. 6I). Yet, for the 168- persistence in the LLNL TSB and TSB succinate conditions, the community structure change was not as significant based on the beta diversity and composition profiles even though persistence was still revealed by sequencing (Fig. 6, Supp. Fig. 24C-D). For 6051a-persistence in the LLNL M9 condition, the alpha diversity and richness metrics quantified a sustainment of diversity and richness in comparison to the reduction caused by the 168-persistence and beta diversity PCA plots and composition profiles revealed minimal community structure change from the 6051a-persistence compared to the control (Fig. 6, Supp. Fig. 19C-D, 24E). Accordingly, this example demonstrated that the *B. subtilis* strains had varying effects on the SynCom structure after persisting which is further supported by the results observed for the 168- persistence in LLNL NB, TSB and TSB succinate (Fig. 6, Supp. Fig. 24C-E).

**Figure 6.**
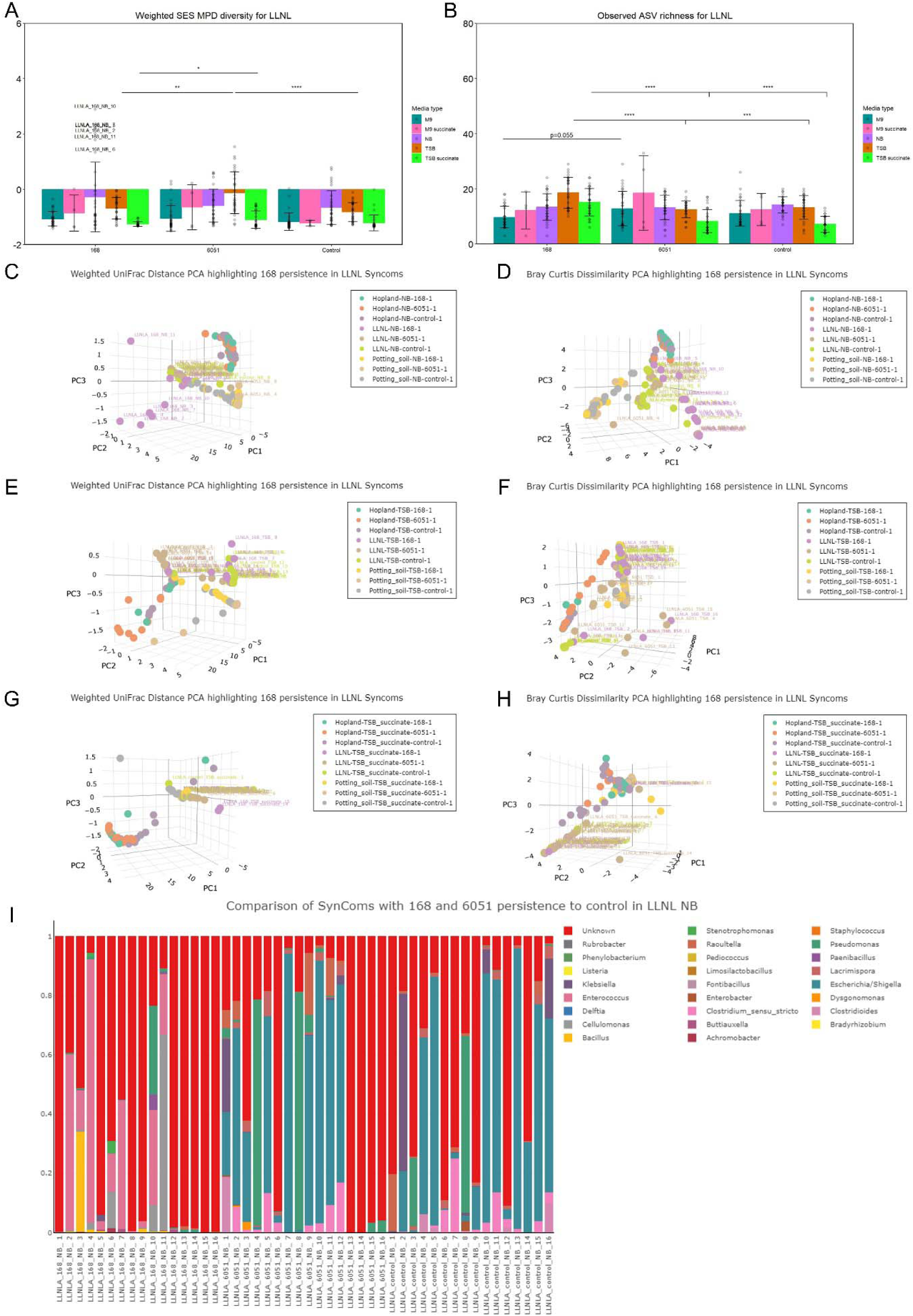
*B. subtilis* 168 strain increased diversity, richness and beta diversity distance of SynComs because of inoculation in LLNL soil. After analyzing all the SynComs (control or base and inoculation conditions) post 16S rRNA sequencing, several metrics were calculated to assess the change in the SynComs after inoculating the SynComs with 168 and 6051a (6051) strains. Standardized effect size (SES) of mean pairwise distance (MPD) phylogenetic diversity is shown for alpha diversity (A) and Observed amplicon sequencing variant (ASV) richness for estimating richness (B) with inter-inoculation statistics shown. For beta diversity, Weighted UniFrac distance (C, E, G) and Bray Curtis Dissimilarity (D, F, H) are visualized as PCA plots with graphs grouped based on media type then comparing soil and inoculation type. Specific labels are placed on the PCA plots to show which SynComs separated from others. A representative composition profile at the genus level (I) is shown to exemplify the impact from the inoculation process. Data for alpha diversity is mean ± SD with each SynCom represented as a point which is up to 32 individual replicates as there were 16 SynComs in each media type of every soil and two separate 96-well plates were sequenced for every condition with the same for PCA plots; *p<0.05, **p<0.01 ***p<0.001, ****p<0.0001.

To show a more holistic hierarchical analysis of the inoculation impact on all conditions, a phylogenetic tree was constructed to compare the tree structure of the control SynComs to all SynComs displayed how the inoculation process disrupted the grouping of the SynComs strictly by soil to more distribution regarding soil, but some grouping based on inoculation condition (Supp. Fig. 25). Lastly, when comparing the *B. subtilis* strain’s effect across the soil and media types, most of the significant changes stemmed from the initial complexity of the control SynComs (i.e., the more diverse the initial SynCom, the more diverse after inoculation) but in some of the conditions with persistence, the diversity and richness that resulted was higher because of the persistence and not dependent on the starting diversity or richness (Supp. Fig. 26). Three-dimensional visuals of beta diversity and selective composition profile visuals were generated to provide further context and comparisons beyond the two-dimensional visuals (GitHub, Code Availability statement).

## Discussion

In this work, we demonstrated a highly parallelized top-down, agnostic approach to cultivation and characterization of SynComs that can yield diverse consortia representing a target soil ecology[51,60]. The compositions of these stable, often complex SynComs varied significantly and stratified primarily based on soil source (i.e. composition of the initial metacommunity prior to selection), as expected, although varying the growth medium led to clear and, in some cases, dramatic compositional variation between cultivated SynComs. The ecological richness observed in SynComs derived from the Hopland and Potting soil samples likely reflects the nutritional density of these agricultural like soils and the relatively extensive biodiversity they support of those environments[61–63]. When evaluating the variance (e.g., composition, diversity and richness) between SynComs, we found that independent samplings of the same source soil (referred to in this report as “biological” replicates) can yield markedly different SynCom structures even when propagated in the same media especially in the more nutritionally dense soils[64], whereas a single soil sampling propagated two or more times independently in the same media (referred to in this report as “technical” replicates) often yields SynComs with similar composition and structure (Supp. Fig. 6-7). These observations underscore several important practical considerations for the isolation and characterization of *in vitro* microbial communities: on the one hand, a single soil source can be a rich source of novel microbial diversity, even when repeatedly sampled, and especially when cultivated in parallel in several divergent conditions; on the other hand, consistency in the profiles of “technical replicate” communities imply reproducibility in community-scale experiments and hint at a deterministic quality to the dynamics of wild communities subjected to ecological selection *in vitro*. Though the source soil provided the reservoir of biodiversity, our results are consistent with prior reports showing that media composition plays an outsize role in shaping community composition[65–67]. We found that cultivation in rich, undefined media generally led to greater diversity and richness relative to minimal, defined media, but PCA indicates that SynComs produced in these varying conditions have non-overlapping compositions, consistent with the notion that cultivation in a variety of media types is advantageous in isolating diverse, divergent consortia, including those containing previously undescribed organisms (Supp. Fig. 6).

Incorporating succinate as an alternative carbon source proved particularly valuable, as we observed increased complexity, particularly in the SynComs derived from agricultural like soils, following addition of succinate to either rich or minimal media, potentially reflecting the typical presence of succinate in root exudates and utilization by a variety of rhizosphere bacteria[52,53]. When comparing the stability of SynComs during a storage and regrowth process, most SynComs exhibited diversity retention with some fluctuations in community structure but overall preservation indicating capacity to reuse SynComs for other purposes if needed. Overall, this top-down approach to SynCom cultivation provided a powerful but straightforward means to generate diverse, stable SynComs derived from a target environment.

We further confirmed that this community cultivation approach could be readily adapted to screen for the presence of exogenously supplied genetically-modified biofertilizer organism to screen for persistence with native consortia passaged over many generations. Heterologous introduction of a gene encoding inducible secreted β-galactosidase in the *B. subtilis* strains provided a valuable screening tool that enabled quantitative, temporal, non-destructive tracking of β-gal activity across many communities in parallel as a proxy for the presence and relative abundance of the introduced *B. subtilis* strain(s), allowing efficient down-selection of target communities for profiling. Although our induction and screening protocols (developed for simplicity and compatibility with automation) could be optimized to improve induction conditions (and therefore detection sensitivity), the calibration curves provided sufficient sensitivity to accurately identify *B. subtilis*-positive SynComs even at relative abundances of ∼1%, along with temporal context for fluctuations in metabolic activity and abundance (Fig. 4, Supp. Fig. 11-12). While β-gal was selected to provide sensitive, low-background detection with minimal potential to perturb community dynamics, alternative screening reporters and designs can be envisioned, those highlighting specific target traits or functions; however limited foreknowledge of the phenotypic landscape of a soil metacommunity can hamper screening success, particularly if cultivate organisms exhibit phenotypes that confound or interfere with screening assays (e.g. fluorescence, bioluminescence, etc.). For our purposes, the ONPG-based screening approach represents a viable proof-of-concept method that can be used to identify target SynComs for cost-effective downstream analysis, such as sequencing (Fig. 4, Supp. Fig. 11-12).

Sequencing experiments generally revealed the presence of the heterologous *B. subtilis* strains that was indicated by β-gal activity (Fig. 4, Supp. Fig. 12-18). Furthermore, observed differences between performance of *B. subtilis* strains utilized may reflect different basal metabolic capabilities of these strains; for example, strain 6051a generally exhibited improved persistence with SynComs in minimal (M9) medium relative to 168, potentially owing to the known tryptophan auxotrophy of the 168 strain[68]. Similarly, the reduced persistence of *B. subtilis* in SynComs cultivated in the presence of succinate may reflect the complex impact on central metabolic flux owing to the presence of organic acids in the growth medium and ease of continued acidification of medium[69,70]. Ultimately, we demonstrated that multiple strains of a representative biofertilizer (*B. subtilis*) organism can persist within top-down cultivated *in vitro* SynComs from multiple soil and media types across a range of community structures (Supp. Fig. 11). Having indication of persistence of *B. subtilis* strains in the sampled SynComs, we investigated the impact of *B. subtilis* incorporation on SynCom structure and stability. In SynComs isolated from Hopland and Potting soil (i.e., the more biodiverse soil types sampled), the inoculation of *B. subtilis* is frequently correlated with reduced community diversity and richness, although a wide array of divergent outcomes were observed, including minimal impact on community composition following *B. subtilis* incorporation (Fig. 5, Supp. Fig. 19-22).

Whereas diversity and richness were increased in Livermore soil-derived SynComs specifically in the conditions where *B. subtilis* persisted (Fig. 6, Supp. Fig. 24). In fact, the presence or absence of *B. subtilis* became a key variable that impacted Livermore soil-derived SynCom structure which could be explained by *B. subtilis* fulfilling an environmental niche in these SynComs. The persistence of *B. subtilis* in SynComs often appeared to correlate with the taxonomic composition of the recipient consortium than the soil source or media type; overrepresented in *B. subtilis-*containing SynComs are PGPR members that have been shown to interact with *Bacillus* such as *Pseudomonas*[71–73], *Klebsiella*[74,75]*, Stenotrophomonas*[76,77] and other similar spore forming bacteria such as *Paenibacillus*[78–80] along with multiple unknown genera. We hierarchically clustered SynComs based on compositional profile and ascertained that the introduction of *B. subtilis* disrupted soil-type-based groupings and introduced some new persistence-based groupings, suggesting that *B. subtilis* incorporation had an impact on community structure overall (Supp. Fig. 25), although, again, a wide array of outcomes complicates the elucidation of any firm trends, underscoring the complexity of microbial community dynamics, even in relatively contrived laboratory conditions. We argue that these divergent outcomes can be leveraged to benefit for the purpose of developing microbial communities containing specific organisms of interest; while far too little data exists to support robust bottom-up design of polymicrobial communities containing target engineered organisms, the process described here affords a means to readily isolate such communities, and indicates that many permutations can be isolated and stably cultivated even from a single soil source.

Overall, the approach described here provides a high-throughput mechanism for cultivating complex microbial communities and screening them for the presence of artificially introduced biofertilizer strains (here, *B. subtilis*). Characterization of the impact of biofertilizer strains on (and compatibility with) native consortia could be further automated and expanded to profile an array of soil types and ecologies to optimize deployment of biofertilizer organisms.

Additionally, this method can be used in a parallel bioprospecting campaign to identify organisms exhibiting desired functionality that could potentially synergize with or even supplant a particular chassis organism. For example, carbon and nitrogen biofertilizer microbes are currently being deployed in field trials without profiling persistence capability (potentially to the detriment of performance) against the native consortia, and with a limited available toolbox of organisms to remediate failures[81–83]. The strengths of the top-down SynCom approach (simplicity in design, broad discovery potential, robustness to complexity), on the other hand, can be leveraged to achieve function- or environment-specific outcomes that can ultimately optimize performance and persistence of target organisms in native ecosystems[78,79].

Moreover, this high-throughput approach could generate data needed for machine learning and artificial intelligence models to better understand community assembly dynamics and collaborative functionality, ultimately enabling predictions that significantly optimize biofertilizer- based technologies, whether globally or for specific soil types and applications.

## Methods

### Generation of top-down SynComs

Top-down SynComs were cultivated following protocols developed previously[51] but with the additions of several soil and media types. Bulk soil from the University of California (UC) Hopland Research and Extension Center (Hopland), in southwestern Mendocino County, CA (39° 00′ 14.6″ N, 123° 05′ 09.1″ W)[55]; the field station exists on territory originally home to the indigenous Pomo Nation, bulk soil from Lawrence Livermore National Laboratory (LLNL) and Potting soil (Home Depot, Kellogg Garden Organics Indoor Potting Mix #3037) were used to create a variety of soils to culture diverse communities. The media types included standard M9 minimal medium (Cold Spring Harbor protocol recipe), nutrient broth (NB; BD Difco, #234000) and tryptic soy broth (TSB; Sigma-Aldrich, #22092). Additionally, these three media types were modified to contain equimolar concentrations of succinate (Sigma-Aldrich, S3674) to glucose with the NB concentration equally that of TSB. NB and TSB containing succinate were adjusted to pH 6.5 to match the pH occurring in M9 with succinate. These six media types were distributed column-wise in the 96-well plates creating 16 replicates of each media type. Five grams of each soil was resuspended in 50 ml of phosphate buffered saline (PBS, pH 7.4) and cycloheximide at 200 µg/ml and incubated at room temperature for 48 h. The large particles were allowed to settle by gravity and 50 µl of the supernatant was distributed into 450 µl of media in each well of a 96-deep well V bottom plate (Opentrons, # 999-00103). This process was repeated with separate five grams of soil from each soil type for three separate 96-well plate replicates (9 plates total). After the initial distribution of the supernatant, the communities were grown in the deep well V bottom plates, after being covered with breathe easy film (USA Scientific, #9123-6100), at 30°C and static (no shaking) conditions for 48 h before the communities were passaged. For passaging, the plates were vortexed (70% strength of Vortex-Genie 2) for 15 seconds then 5 µl is passaged into 495 µl of fresh media after pipetting up and down several times. This process was repeated 10 times to establish stable communities[60] then the communities were stored at -80°C after combining 50 µl of each community with 50 µl of 50% glycerol. When the communities were reanimated, the entire mixture of the stored communities was transferred into 400 µl of fresh media then grown for 72 h during the initial reanimation before passaging. After the next passage, the communities were restored for potential use. For sequencing, the communities were sequenced after the first and second reanimations to compare stability across freeze-thaw cycles. Additionally, the communities were grown for an additional two passages post reanimation before sequencing to have a direct comparison to the inoculated communities described below as the screening assay for persistence revealed some fluctuations in the community dynamics in the initial reanimation and first passage but a re-establishment of stability by the second passage post reanimation.

### *B. subtilis* 168 and 6051a constructs

Constructs were inserted into the genome of *B. subtilis* 168 and 6051a (ATCC #23857 and *Bacillus subtilis* (Ehrenberg) Cohn (ATCC 6051a), ATCC, Manassas, VA, USA) at the *amyE* locus using a homologous recombination plasmid (pDR111[84], a gift from Dr. Lee Kroos). The pDR111 plasmid was transformed into *B. subtilis* 168 and 6051a using a natural competence protocol and constructs were selected by spectinomycin then confirmed by PCR amplification out of the genome[38]. The xylose-inducible system[57,58] was amplified from *B. subtilis* 168 strain genome. The xylose promoter and regulator were cloned into the pDR111 plasmid to replace the Phyper-spank promoter and LacI regulator using Gibson assembly. *B. subtilis* 168 and 6051a were designed to secrete β-gal to the supernatant through the twin-arginine translocation (Tat) pathway by fusing the PhoD signal peptide[59] to *lacZ* after *lacZ* was amplified from the pST5832 plasmid (a gift from Carolyn Bertozzi & Jessica Seeliger, Addgene plasmid #36256) then the fusion was cloned into the NheI restriction site present in the pDR111 xylose plasmid using Gibson assembly.

### Growth conditions for engineered *B. subtilis* 168 and 6051a

*B. subtilis* 168 and 6051a were grown in Luria-Bertani Miller broth (LB) with spectinomycin (100 µg/ml) before growth curves, β-gal induction and calibration experiments and adding to SynComs. The overnight cultures were grown for 24 h at 30°C and 250 RPM.

### Growth curve experiments for engineered *B. subtilis* 168 and 6051a

*B. subtilis* 168 and 6051a strains expressing xylose-inducible β-gal were grown as described above in triplicate (n = 3). All cultures were spun down and normalized to OD_600_ = 1 in PBS then a 1:100 was performed into every media type from SynCom studies in a 96 well plate (ThermoFisher Scientific, #267544) for a total of 12 replicates per media type and strain (n = 3 biological, n = 4 technical). Cultures were grown in a BioTek Synergy HTX plate reader at 30°C and 425 CPM (fast orbital shaking) with OD_600_ measurements taken every 30 min using Gen 53.11.19 software. Measurements were performed for 40 h. All replicates were plotted to visualize differences in growth rates between strains as mean ± SEM.

### β-gal induction and calibration experiments for engineered *B. subtilis* 168 and 6051a

Engineered *B. subtilis* 168 and 6051a were grown as described above before being prepared for induction experiments. For the induction experiments, the cultures for both strains were spun down and resuspended in PBS in triplicate before being distributed into LB in 96-well assay plates (Fisher Scientific, #07-200-656). Each biological replicate was serially diluted in three columns (n = 3 biological and n = 3 technical) starting at 100 million cells then performing 1:2 dilutions in 100 µl down the rows in the 96-well plates. Replicate 96-well plates were used to duplicate the serial dilutions without xylose in one plate and the other plate contained 0.5% D-xylose (Sigma Aldrich, #W360600) in every well. After dilutions, the assay plates were incubated at 30°C and static conditions for 2 h then ONPG was added. ONPG (ThermoFisher Scientific, #34055) was aliquoted into every well at 1 mg then absorbance was read immediately after at 420 nm on a BioTek Synergy H1 plate reader using Gen 5 1.11 software. Incubation was continued at 30°C for 1 h then absorbance was read at 420 nm again to determine the change in absorbance based on cell number. For calibration experiments, engineered *B. subtilis* 168 and 6051a were grown as described above in triplicate (n = 3) then were spun down and resuspended in PBS at OD_600_ = 1 before being subcultured into the six media types in 96-deep well plates used in the SynCom experiments to replicate the same conditions that would be used for inoculation into SynComs. Once normalized to OD_600_ = 1, the strains were diluted 1:100 in 1 ml of TSB while for the other 5 media conditions the strains were diluted 1:4 because of results observed in the growth curves and all the dilutions were done in quadruplicate (n = 4) for each biological replicate of each strain in every media condition. Afterwards, the conditions were then grown at 30°C at static conditions for 48 h in the 96-well deep well plates before proceeding to serial dilutions for the calibration curves. The process used in the induction curves was reproduced under conditions with 0.5% D-xylose with modifications to the starting number of cells because of the limited growth of the strains in certain media types. Accordingly, 100 million cells was used as the starting point in TSB while for M9 and M9 succinate 20 million cells was used and for the other three media types (NB, NB succinate, TSB succinate) 50 million cells was used. The rest of the protocol from the induction was followed exactly. To create the best fit polynomial for use during the inoculation into SynComs to generate a general sense of *Bacillus* cell number because of phenotype and display the formulas for the curves, R functions including stat_poly_line and stat_poly_eq were used and second-degree polynomial was found to be ideal.

### Inoculation of engineered *B. subtilis* 168 and 6051a into top-down SynComs

Stable SynComs generated above were reanimated and grown for 72 h then the SynComs were passaged into separate parallel plates to maintain the base SynComs as background in the ONPG assays described below and to start inoculation into SynComs. *B. subtilis* 168 and 6051a were grown as described above before preparation for separately adding to all base SynComs (three soil types with three separate five gram replicates equaling 9 plates total for each strain). The strains were spun down and resuspended in PBS at a normalized of OD_600_ = 1. The stable SynComs were passaged after the 72 h of growth then 5 µl of the normalized strains was distributed into every well. The plates with the addition the engineered *B. subtilis* 168 and 6051a were passaged just as the stable SynComs above except the number of passages was modified. The passages were determined from the ONPG tracking experiment described below and it was observed that five passages provided multiple passages of stability leading up to a freeze thaw cycle where it was observed that an additional two passages post the reanimation growth resulted in stability of phenotype again. Therefore, five passages were used for persistence testing after initial addition then two passages were used after the initial reanimation cycle following the freeze-thaw before proceeding to sequencing.

### ONPG tracking of engineered *B. subtilis* 168 and 6051a persistence

To track the engineered *B. subtilis* 168 and 6051a strains interaction with the native microbial consortia and down select communities to sequence, ONPG assays following the protocols established above (β-gal induction and calibration) were performed at every passage. After all the base, stable SynComs were grown up and split into separate plates to prepare for inoculation mentioned above, both the stable SynComs and the inoculation conditions were used in the ONPG assays to determine background from the starting communities and changes resulting from persistence. The base SynComs were tracked for multiple passages to establish background while the inoculation conditions were tracked for six passages post initial addition of the strains (P0) and after the sixth passage (P5) the inoculated communities went through a freeze-thaw cycle. This process was determined by observing stability of signal for multiple passages. After the freeze-thaw cycle, the inoculated communities were passaged and tracked using the ONPG assays until stability of signal was reestablished (P8). For every ONPG screen, 200 µl of the SynComs were aliquoted into the 96-well assay plates then the assays developed in the β-gal induction and calibration curves was followed exactly. The change in absorbance in the base SynComs (background) was subtracted from the change in absorbance in the inoculated communities to determine the true change in absorbance. The calibration curves established for every media type for both strains were then used following the quadratic equation to estimate the relative abundance of the engineered *B. subtilis* strains over time. These assays were used to select which replicate plates and media conditions could be best for sequencing and the ONPG results for the 1200 communities selected were used to determine effectiveness of the assay due to cross-validation with sequencing. Ordinary one-way ANOVAs were used initially to determine significance between visualized conditions and Bartlett’s test was applied to assess equality of variances. We found that the equality of variances was not upheld so we utilized Brown-Forsythe and Welch’s one-way ANOVA with Dunnett’s T3 post-hoc test to determine significance.

### Genomic DNA preparation for sequencing

Extraction of gDNA from the communities was performed using a modification from protocols developed previously[85]. A 100 µl aliquot of the communities was combined with 100 µl of 1% IGEPAL CA-630 (Sigma Aldrich, #I8896) in a 96-well rigid, semi-skirted PCR plate (ThermoFisher Scientific, #AB0990) then the previous protocols including three freeze-thaw cycles and adding Proteinase K at 100 µg/ml then incubating at 60°C for 1 h were followed (Method 3)[85]. Following the above lysis protocol, the gDNA was cleaned following Illumina’s 16S metagenomic library preparation protocol (PCR cleanup 2). AMPure XP beads (Beckman Coulter, #A63881) were vortexed for 30 seconds then 112 µl of beads were added to 112 µl of the lysed communities in a 96-well rigid, semi-skirted PCR plate then the rest of the protocol was followed as described in Illumina’s protocol. Once the gDNA was purified, the gDNA was quantified using a Quant-iT dsDNA high sensitivity assay kit (ThermoFisher Scientific, #Q33120) before submitting to SeqCenter for sequencing.

### 16S rRNA amplicon sequencing

All samples (1200) and the community standard (Zymo Research, #D6300) were submitted to SeqCenter for sequencing. The samples were sequenced for 50k reads and every sample reached >80k reads with most samples achieving >100k reads. Samples were prepared using Zymo Research’s Quick-16S kit with phased primers targeting the V3/V4 regions of the 16S gene with the following sequences: CCTACGGGDGGCWGCAGCCTAYGGGGYGCWGCAG (forward) and GACTACHVGGGTATCTAATCC (reverse). Following clean up and normalization, samples were sequenced on a P1 600cyc NextSeq2000 Flowcell to generate 2x301bp paired end (PE) reads.

### 16S rRNA amplicon analysis

Primer sequences were 5’ trimmed using cutadapt v4.6[86]. Trimmed reads were further filtered for quality with DADA2 v1.28.0[87] (parameters maxN = 0, maxEE = c(2, 2), truncQ = 2) and the forward and reverse reads were right trimmed to 270 and 200, respectively. Chimeric sequences were predicted *de novo* and removed with the DADA2 removeBimeraDenovo() function in “consensus” mode. The resulting amplicon sequence variants (ASVs) were assigned taxonomy with RDP training set 18[88], were aligned with mafft v7.505[89] and were used to build a phylogenetic tree with fasttree 2.1.11[90] followed by midpoint rooting with phytools v2.3- 0[91]. Once a phyloseq object was created, R packages such as phyloseq and picante were used to analyze the results in the object then create visualizations accordingly based on further packages described in statistical analysis and data visualization below. For the control top-down SynComs and inoculated SynComs analysis, Ordinary one-way ANOVAs were used initially to determine significance between visualized conditions and Bartlett’s test was applied to assess equality of variances. We found that in many of the conditions the equality of variances was not upheld so we then utilized Brown-Forsythe and Welch’s one-way ANOVA with either Games-Howell’s post-hoc test (n > 50 in soil only grouping) or Dunnett’s T3 post-hoc test (n < 50 in all other conditions) to determine significance. Additionally, when the conditions were broken down by soil and media type and one condition did not have enough samples (n > 2) to be included in the appropriate ANOVA (e.g., M9 succinate), then unpaired two-tailed T tests were utilized initially, and F test was applied to assess equality of variances. We found that in many of the conditions the equality of variances was not upheld so we then utilized Welch’s unpaired two-tailed T tests to determine significance.

### Statistical analysis and data visualization

Statistical analyses were performed using Prism software (10.2.3, GraphPad Inc., La Jolla, CA). Statistical tests are identified for each method in the sections above. Data was considered significant if *p*<0.05. Plotting was performed using R version 4.4.0 with the following packages: phyloseq 1.48.0[92], picante 1.8.2[93], RColorBrewer 1.1-3, GUniFrac 1.8[94], viridis 0.6.5[95], ape 5.8[96], plotly 4.10.4, shiny 1.8.1.1, shinyjs 2.1.0, htmlwidgets 1.6.4, highcharter 0.9.4, ggplot2 3.5.1, ggsignif 0.6.4[97] and ggbreak 3.3.2[98].

## Data Availability statement

All raw data, engineered *B. subtilis* constructs and SynComs will be made available upon request by the corresponding authors. Raw data from all sequencing is uploaded to NCBI SRA as BioProject PRJNA1171870.

## Code Availability statement

All R scripts were written with a general format appropriate for the openly available, established packages mentioned above and can be made available on request. Code for creating 3D visualizations is available on GitHub (https://github.com/madsen16/Synthetic-communities).

## Acknowledgements

The authors would like to thank Dr. Lee Kroos for *B. subtilis* plasmid pDR111. The authors would also like to thank SeqCenter for efficient and cost-effective services. The authors would like to thank Janelle Cataldo from Lawrence Livermore National Laboratory for graphic design of Figure 1. The authors would like to acknowledge the Laboratory Directed Research and Development Program at Lawrence Livermore National Laboratory for funding this work (23-LW-014). This work was performed under the auspices of the DOE by Lawrence Livermore National Laboratory under Contract DEAC52-07NA27344 (LLNL-JRNL-2002743-DRAFT). Supplementary Figure 1-2 were created using BioRender.com.

## Author contributions

Dr. Cody S. Madsen conceptualized the methods for inoculating *B. subtilis* into SynComs, developed all the engineered strains, performed all experiments, jointly developed and analyzed all data to make figures and was one of the primary authors of the manuscript. Dr. Jeffrey A. Kimbrel analyzed all sequencing data, aided in developing figures and was one of the primary authors of the manuscript. Dr. Patrick Diep supported preparation of gDNA for sequencing of the 1200 SynComs and significantly contributed to the writing and editing of this manuscript. Dr. Dante P. Ricci conceptualized the SynCom cultivation process and significantly contributed to the writing and editing of this manuscript.

## References

1. Chakraborty T, Akhtar N. Biofertilizers: prospects and challenges for future. Biofertilizers: study and impact 2021:575–90.

2. Rai PK, Rai A, Sharma NK et al. Limitations of biofertilizers and their revitalization through nanotechnology. J Clean Prod 2023:138194.

3. Mącik M, Gryta A, Frąc M. Biofertilizers in agriculture: An overview on concepts, strategies and effects on soil microorganisms. Advances in agronomy 2020;162:31–87.

4. Mahmud AA, Upadhyay SK, Srivastava AK et al. Biofertilizers: A Nexus between soil fertility and crop productivity under abiotic stress. Current Research in Environmental Sustainability 2021;3:100063.

5. Kumar S, Sindhu SS, Kumar R. Biofertilizers: An ecofriendly technology for nutrient recycling and environmental sustainability. Curr Res Microb Sci 2022;3:100094.

6. Compant S, Cassan F, Kostić T et al. Harnessing the plant microbiome for sustainable crop production. Nat Rev Microbiol 2024:1–15.

7. Bloch SE, Clark R, Gottlieb SS et al. Biological nitrogen fixation in maize: optimizing nitrogenase expression in a root-associated diazotroph. J Exp Bot 2020;71:4591–603.

8. Wen A, Havens KL, Bloch SE et al. Enabling Biological Nitrogen Fixation for Cereal Crops in Fertilized Fields. ACS Synth Biol 2021;10:3264–77.

9. Ke X, Feng S, Wang J et al. Effect of inoculation with nitrogen-fixing bacterium Pseudomonas stutzeri A1501 on maize plant growth and the microbiome indigenous to the rhizosphere. Syst Appl Microbiol 2019;42:248–60.

10. Schnabel T, Sattely E. Improved Stability of Engineered Ammonia Production in the Plant-Symbiont Azospirillum brasilense. ACS Synth Biol 2021;10:2982–96.

11. Schnabel T, Sattely E. Engineering posttranslational regulation of glutamine synthetase for controllable ammonia production in the plant symbiont Azospirillum brasilense. Appl Environ Microbiol 2021;87:e00582–21.

12. Connolly A. Are Microbes The Future Of Fertilizer? Forbes 2023.

13. Aranda-Díaz A, Ng KM, Thomsen T et al. Establishment and characterization of stable, diverse, fecal-derived in vitro microbial communities that model the intestinal microbiota. Cell Host Microbe 2022;30:260–72.

14. Bittleston LS, Gralka M, Leventhal GE et al. Context-dependent dynamics lead to the assembly of functionally distinct microbial communities. Nat Commun 2020;11:1440.

15. van Leeuwen PT, Brul S, Zhang J et al. Synthetic microbial communities (SynComs) of the human gut: design, assembly, and applications. FEMS Microbiol Rev 2023;47:fuad012.

16. Pascual-García A, Bonhoeffer S, Bell T. Metabolically cohesive microbial consortia and ecosystem functioning. Philosophical Transactions of the Royal Society B 2020;375:20190245.

17. George AB, Korolev KS. Ecological landscapes guide the assembly of optimal microbial communities. PLoS Comput Biol 2023;19:e1010570.

18. Marín O, González B, Poupin MJ. From Microbial Dynamics to Functionality in the Rhizosphere: A Systematic Review of the Opportunities With Synthetic Microbial Communities. Front Plant Sci 2021;12:650609.

19. Liu Y-X, Qin Y, Bai Y. Reductionist synthetic community approaches in root microbiome research. Curr Opin Microbiol 2019;49:97–102.

20. Vorholt JA, Vogel C, Carlström CI et al. Establishing causality: opportunities of synthetic communities for plant microbiome research. Cell Host Microbe 2017;22:142–55.

21. Ishizawa H, Tada M, Kuroda M et al. Synthetic bacterial community of duckweed: a simple and stable system to study plant-microbe interactions. Microbes Environ 2020;35:ME20112.

22. Bodenhausen N, Bortfeld-Miller M, Ackermann M et al. A synthetic community approach reveals plant genotypes affecting the phyllosphere microbiota. PLoS Genet 2014;10:e1004283.

23. Trimmer M, Grey J, Heppell CM et al. River bed carbon and nitrogen cycling: state of play and some new directions. Science of the total environment 2012;434:143–58.

24. Martínez-Espinosa RM. Microorganisms and their metabolic capabilities in the context of the biogeochemical nitrogen cycle at extreme environments. Int J Mol Sci 2020;21:4228.

25. de Jesús Astacio LM, Prabhakara KH, Li Z et al. Closed microbial communities self-organize to persistently cycle carbon. Proceedings of the National Academy of Sciences 2021;118:e2013564118.

26. Thamdrup B. New pathways and processes in the global nitrogen cycle. Annu Rev Ecol Evol Syst 2012;43:407–28.

27. Barnawal D, Maji D, Bharti N et al. ACC deaminase-containing Bacillus subtilis reduces stress ethylene-induced damage and improves mycorrhizal colonization and rhizobial nodulation in Trigonella foenum-graecum under drought stress. J Plant Growth Regul 2013;32:809–22.

28. Misra S, Chauhan PS. ACC deaminase-producing rhizosphere competent Bacillus spp. mitigate salt stress and promote Zea mays growth by modulating ethylene metabolism. 3 Biotech 2020;**10**:1–14.

29. Hecker M, Völker UBT-A in MP. General stress response of Bacillus subtilis and other bacteria. Vol 44. Academic Press, 2001, 35–91.

30. Völker U, Maul B, Hecker M. Expression of the sigmaB-dependent general stress regulon confers multiple stress resistance in Bacillus subtilis. J Bacteriol 1999;181:3942–8.

31. Hashem A, Tabassum B, Abd_Allah EF. Bacillus subtilis: A plant-growth promoting rhizobacterium that also impacts biotic stress. Saudi J Biol Sci 2019;26:1291–7.

32. Vocciante M, Grifoni M, Fusini D et al. The Role of Plant Growth-Promoting Rhizobacteria (PGPR) in Mitigating Plant’s Environmental Stresses. Applied Sciences 2022;12, DOI: 10.3390/app12031231.

33. de Lima BC, Moro AL, Santos ACP et al. Bacillus subtilis ameliorates water stress tolerance in maize and common bean. J Plant Interact 2019;14:432–9.

34. Elshaghabee FMF, Rokana N, Gulhane RD et al. Bacillus As Potential Probiotics: Status, Concerns, and Future Perspectives. Front Microbiol 2017;8:1490.

35. Liu Y, Liu L, Li J et al. Synthetic Biology Toolbox and Chassis Development in Bacillus subtilis. Trends Biotechnol 2019;37:548–62.

36. Jakutyte-Giraitiene L, Gasiunas G. Design of a CRISPR-Cas system to increase resistance of Bacillus subtilis to bacteriophage SPP1. J Ind Microbiol Biotechnol 2016;43:1183–8.

37. Zhang K, Duan X, Wu J. Multigene disruption in undomesticated Bacillus subtilis ATCC 6051a using the CRISPR/Cas9 system. Sci Rep 2016;6:27943.

38. Harwood CR, Cutting SM. Molecular biological methods for Bacillus. 1990.

39. Rokop ME, Auchtung JM, Grossman AD. Control of DNA replication initiation by recruitment of an essential initiation protein to the membrane of Bacillus subtilis. Mol Microbiol 2004, DOI: 10.1111/j.1365-2958.2004.04091.x.

40. Loewus FA, Kelly S, Neufeld EF. Metabolism of myo-inositol in plants: conversion to pectin, hemicellulose, D-xylose, and sugar acids. Proc Natl Acad Sci U S A 1962;48:421.

41. Ankel H, Feingold DS. Biosynthesis of uridine diphosphate D-xylose. I. Uridine diphosphate glucuronate carboxy-lyase of wheat germ. Biochemistry 1965;4:2468–75.

42. Baron D, Wellmann E, Grisebach H. Purification and properties of an enzyme from cell suspension cultures of parsley catalyzing the synthesis of UDP-apiose and UDP-D-xylose from UDP-D-glucuronic acid. Biochim Biophys Acta 1972;258:310–8.

43. Vavrová Ľ, Muchová K, Barák I. Comparison of different Bacillus subtilis expression systems. Res Microbiol 2010, DOI: 10.1016/j.resmic.2010.09.004.

44. Gärtner D, Geissendörfer M, Hillen W. Expression of the Bacillus subtilis xyl operon is repressed at the level of transcription and is induced by xylose. J Bacteriol 1988, DOI: 10.1128/jb.170.7.3102-3109.1988.

45. Sun T, Altenbuchner J. Characterization of a mannose utilization system in bacillus subtilis. J Bacteriol 2010, DOI: 10.1128/JB.01673-09.

46. Promchai R, Promdonkoy B, Tanapongpipat S et al. A novel salt-inducible vector for efficient expression and secretion of heterologous proteins in Bacillus subtilis. J Biotechnol 2016;222:86–93.

47. Piraner DI, Abedi MH, Moser BA et al. Tunable thermal bioswitches for in vivo control of microbial therapeutics. Nat Chem Biol 2017;13:75–80.

48. Shayanthan A, Ordoñez PAC, Oresnik IJ. The role of synthetic microbial communities (SynCom) in sustainable agriculture. Frontiers in Agronomy 2022;4:896307.

49. Khan ST. Consortia-based microbial inoculants for sustaining agricultural activities. Applied Soil Ecology 2022;176:104503.

50. Pradhan S, Tyagi R, Sharma S. Combating biotic stresses in plants by synthetic microbial communities: Principles, applications and challenges. J Appl Microbiol 2022;133:2742–59.

51. Estrela S, Sanchez-Gorostiaga A, Vila JCC et al. Nutrient dominance governs the assembly of microbial communities in mixed nutrient environments. Kost C, Schuman MC, Waschina S, et al. (eds.). Elife 2021;**10**:e65948.

52. Franzino T, Boubakri H, Cernava T et al. Implications of carbon catabolite repression for plant–microbe interactions. Plant Commun 2022;3.

53. Udvardi M, Poole PS. Transport and metabolism in legume-rhizobia symbioses. Annu Rev Plant Biol 2013;64:781–805.

54. Foley MM, Blazewicz SJ, McFarlane KJ et al. Active populations and growth of soil microorganisms are framed by mean annual precipitation in three California annual grasslands. Soil Biol Biochem 2023;177:108886.

55. Fossum C, Estera-Molina KY, Yuan M et al. Belowground allocation and dynamics of recently fixed plant carbon in a California annual grassland. Soil Biol Biochem 2022;165:108519.

56. Chen J, Zhu Y, Fu G et al. High-level intra-and extra-cellular production of D-psicose 3-epimerase via a modified xylose-inducible expression system in Bacillus subtilis. J Ind Microbiol Biotechnol 2016;43:1577–91.

57. Kim L, Mogk A, Schumann W. A xylose-inducible Bacillus subtilis integration vector and its application. Gene 1996;181:71–6.

58. Vavrová Ľ, Muchová K, Barák I. Comparison of different Bacillus subtilis expression systems. Res Microbiol 2010;161:791–7.

59. Tjalsma H, Bolhuis A, Jongbloed JDH et al. Signal Peptide-Dependent Protein Transport in Bacillus subtilis: a Genome-Based Survey of the Secretome. Microbiology and Molecular Biology Reviews 2000, DOI: 10.1128/mmbr.64.3.515-547.2000.

60. Chang C-Y, Osborne ML, Bajic D et al. Artificially selecting bacterial communities using propagule strategies. Evolution (N Y) 2020;74:2392–403.

61. Shi S, Nuccio E, Herman DJ et al. Successional trajectories of rhizosphere bacterial communities over consecutive seasons. mBio 2015;6:10–1128.

62. Blazewicz SJ, Hungate BA, Koch BJ et al. Taxon-specific microbial growth and mortality patterns reveal distinct temporal population responses to rewetting in a California grassland soil. ISME J 2020;14:1520–32.

63. Greenlon A, Sieradzki E, Zablocki O et al. Quantitative stable-isotope probing (qSIP) with metagenomics links microbial physiology and activity to soil moisture in Mediterranean-climate grassland ecosystems. mSystems 2022;7:e00417-22.

64. Nuccio EE, Starr E, Karaoz U et al. Niche differentiation is spatially and temporally regulated in the rhizosphere. ISME J 2020;14:999–1014.

65. Eevers N, Gielen M, Sánchez-López A et al. Optimization of isolation and cultivation of bacterial endophytes through addition of plant extract to nutrient media. Microb Biotechnol 2015;8:707–15.

66. Kurm V, Van Der Putten WH, Hol WHG. Cultivation-success of rare soil bacteria is not influenced by incubation time and growth medium. PLoS One 2019;14:e0210073.

67. Vieira FCS, Nahas E. Comparison of microbial numbers in soils by using various culture media and temperatures. Microbiol Res 2005;160:197–202.

68. Zeigler DR, Prágai Z, Rodriguez S et al. The origins of 168, W23, and other Bacillus subtilis legacy strains. J Bacteriol 2008, DOI: 10.1128/JB.00722-08.

69. Schilling O, Frick O, Herzberg C et al. Transcriptional and metabolic responses of Bacillus subtilis to the availability of organic acids: transcription regulation is important but not sufficient to account for metabolic adaptation. Appl Environ Microbiol 2007;73:499–507.

70. Singh KD, Schmalisch MH, Stullke J et al. Carbon catabolite repression in Bacillus subtilis: quantitative analysis of repression exerted by different carbon sources. J Bacteriol 2008;190:7275–84.

71. Comeau D, Balthazar C, Novinscak A et al. Interactions between Bacillus spp., Pseudomonas spp. and Cannabis sativa promote plant growth. Front Microbiol 2021;12:715758.

72. Lyng M, Kovács ÁT. Frenemies of the soil: Bacillus and Pseudomonas interspecies interactions. Trends Microbiol 2023;31:845–57.

73. Angelina E, Papatheodorou EM, Demirtzoglou T, et al. Effects of Bacillus subtilis and Pseudomonas fluorescens inoculation on attributes of the lettuce (Lactuca sativa L.) soil rhizosphere microbial community: The role of the management system. Agronomy 2020;10:1428.

74. Shukla N, Singh D, Tripathi A et al. Synergism of endophytic Bacillus subtilis and Klebsiella aerogenes modulates plant growth and bacoside biosynthesis in Bacopa monnieri. Front Plant Sci 2022;13:896856.

75. Biswas S, Philip I, Jayaram S et al. Endophytic bacteria Klebsiella spp. and Bacillus spp. from Alternanthera philoxeroides in Madiwala Lake exhibit additive plant growth-promoting and biocontrol activities. Journal of Genetic Engineering and Biotechnology 2023;21:153.

76. Das PP, Singh KRB, Nagpure G et al. Plant-soil-microbes: A tripartite interaction for nutrient acquisition and better plant growth for sustainable agricultural practices. Environ Res 2022;214:113821.

77. Ercole TG, Kava VM, Aluizio R et al. Co-inoculation of Bacillus velezensis and Stenotrophomonas maltophilia strains improves growth and salinity tolerance in maize (Zea mays L.). Rhizosphere 2023;27:100752.

78. Shank EA, Klepac-Ceraj V, Collado-Torres L et al. Interspecies interactions that result in Bacillus subtilis forming biofilms are mediated mainly by members of its own genus. Proceedings of the National Academy of Sciences 2011;108:E1236–43.

79. Grady EN, MacDonald J, Liu L et al. Current knowledge and perspectives of Paenibacillus: a review. Microb Cell Fact 2016;15:1–18.

80. Govindasamy V, Senthilkumar M, Magheshwaran V, et al. Bacillus and Paenibacillus spp.: potential PGPR for sustainable agriculture. Plant growth and health promoting bacteria 2011:333–64.

81. O’Callaghan M, Ballard RA, Wright D. Soil microbial inoculants for sustainable agriculture: Limitations and opportunities. Soil Use Manag 2022;38:1340–69.

82. Sepúlveda E, Diyarza-Sandoval NA, Guevara-Avendaño E et al. Plant growth-promoting microorganisms from native plants: an untapped resource of biocontrol and biofertilizer agents. Biocontrol Agents for Improved Agriculture. Elsevier, 2024, 29–66.

83. Mikiciuk G, Miller T, Kisiel A et al. Harnessing Beneficial Microbes for Drought Tolerance: A Review of Ecological and Agricultural Innovations. Agriculture 2024;14:2228.

84. Rokop ME, Auchtung JM, Grossman AD. Control of DNA replication initiation by recruitment of an essential initiation protein to the membrane of Bacillus subtilis. Mol Microbiol 2004, DOI: 10.1111/j.1365-2958.2004.04091.x.

85. Song F, Kuehl J V, Chandran A et al. A simple, cost-effective, and automation-friendly direct PCR approach for bacterial community analysis. mSystems 2021;6:10–1128.

86. Martin M. Cutadapt removes adapter sequences from high-throughput sequencing reads. EMBnet journal 2011;17:10–2.

87. Callahan BJ, McMurdie PJ, Rosen MJ et al. DADA2: High-resolution sample inference from Illumina amplicon data. Nat Methods 2016;13:581–3.

88. Wang Q, Cole JR. Updated RDP taxonomy and RDP Classifier for more accurate taxonomic classification. Microbiol Resour Announc 2024;13:e01063–23.

89. Katoh K, Standley DM. MAFFT multiple sequence alignment software version 7: improvements in performance and usability. Mol Biol Evol 2013;30:772–80.

90. Price MN, Dehal PS, Arkin AP. FastTree 2–approximately maximum-likelihood trees for large alignments. PLoS One 2010;5:e9490.

91. 91. Revell LJ. phytools 2.0: an updated R ecosystem for phylogenetic comparative methods (and other things). PeerJ 2024;**12**:e16505.

92. McMurdie PJ, Holmes S. phyloseq: an R package for reproducible interactive analysis and graphics of microbiome census data. PLoS One 2013;8:e61217.

93. Kembel SW, Cowan PD, Helmus MR et al. Picante: R tools for integrating phylogenies and ecology. Bioinformatics 2010;26:1463–4.

94. Chen J, Zhang X, Yang L. GUniFrac: generalized UniFrac distances, distance-based multivariate methods and feature-based univariate methods for microbiome data analysis. R package version 2022;1.

95. Garnier S, Ross N, Rudis B et al. Package ‘viridis.’ 2024.

96. Paradis E, Schliep K. ape 5.0: an environment for modern phylogenetics and evolutionary analyses in R. Bioinformatics 2019;35:526–8.

97. Ahlmann-Eltze C, Patil I. ggsignif: R Package for Displaying Significance Brackets for’ggplot2’. 2021.

98. Xu S, Chen M, Feng T et al. Use ggbreak to effectively utilize plotting space to deal with large datasets and outliers. Front Genet 2021;12:774846.

