## Supplemental figures for "Synthetic communities as a model for determining interactions between a biofertilizer chassis organism and native microbial consortia"

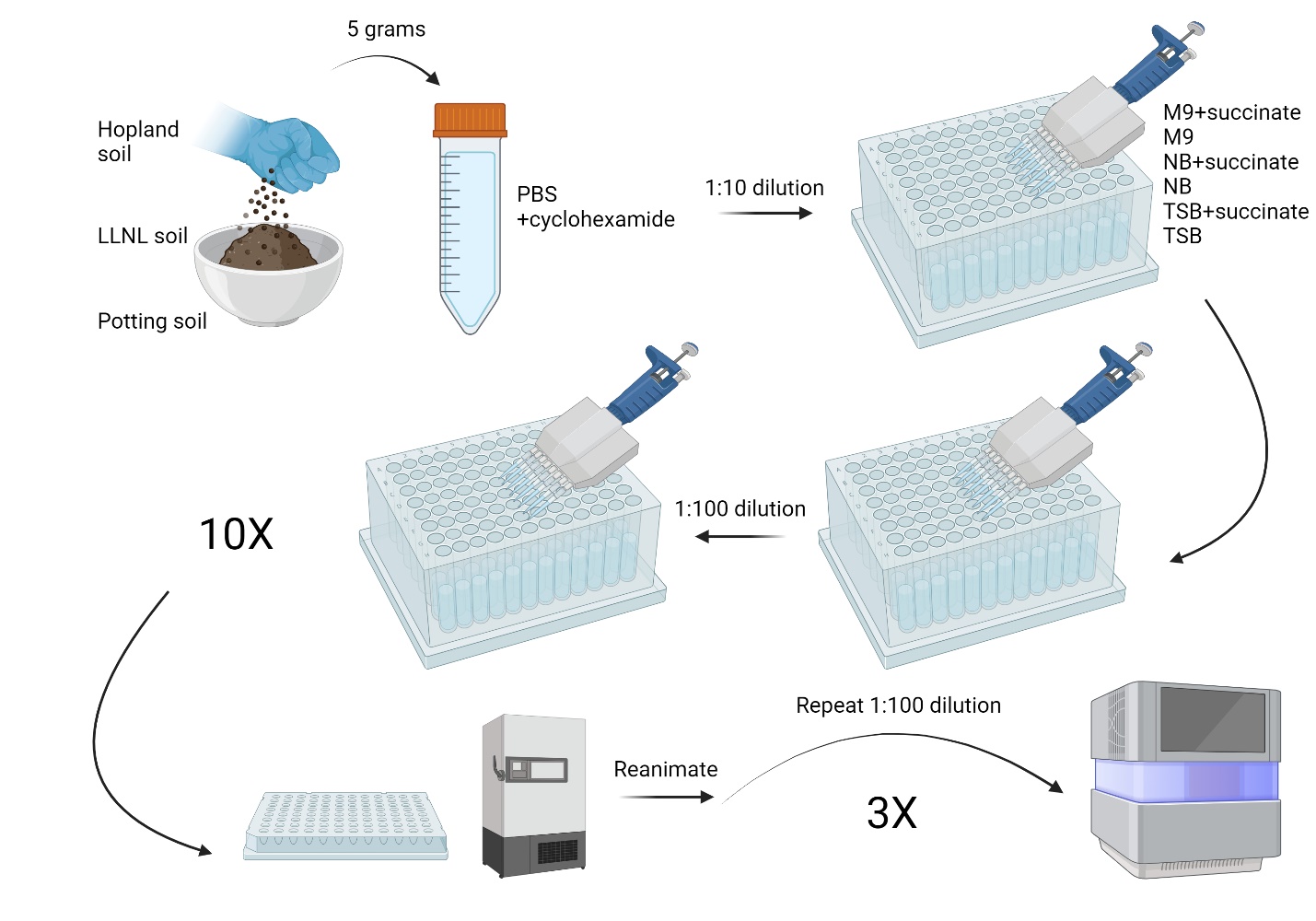

**Figure 1**. **Methodology for cultivating top-down SynComs**

Five grams of soil from Hopland (northern California pasture), Lawrence Livermore National Laboratory and Potting soil (organic indoor potting mix) was resuspended in phosphate buffered saline (PBS) and cycloheximide before the supernatant from the mixture was distributed into six media types in 96 deep-well blocks at a 1:10 dilution. After growing for an initial 48 hours, the cultures were passaged using a 1:100 dilution into fresh media which was then repeated every 48 hours for 10 passages before storing the SynComs. When the SynComs were utilized for inoculation or were sequenced, the SynComs were reanimated and passaged as before for an additional 3 times before preparation for sequencing.

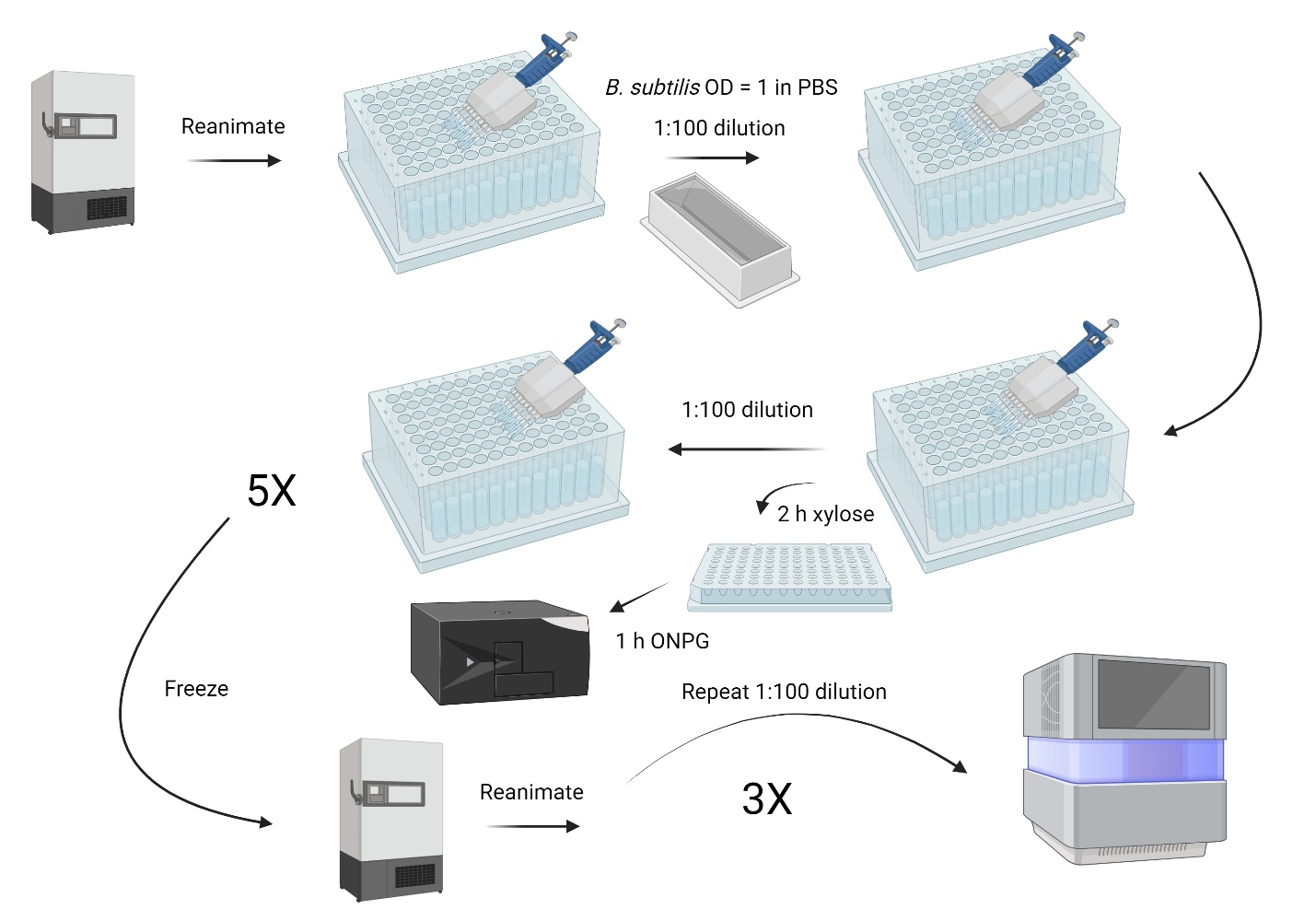

**Figure 2**. **Process for inoculating and screening for *B. subtilis* in established top-down SynComs**

After the established SynComs were reanimated, the *B. subtilis* strains (168 and 6051a) were inoculated into the SynComs by normalizing the strains to OD_600_ = 1 in PBS then adding 5 µl (same amount as SynComs being passaged) to each of the SynComs after the passaging for the SynComs was completed. After the initial 48 hours of outgrowth and the next passage was started, aliquots from the SynComs were added to assay plates to begin screening for presence of *B. subtilis* by inducing secretion of β-gal with xylose for 2 hours. After the 2 hours, ONPG was added to begin colorimetric change for the screening. This process was repeated for 5 additional passages before SynComs were frozen. The SynComs were reanimated to assess potential persistence through a freeze-thaw cycle then 3 additional passages were performed with additional screening before communities were prepared for sequencing.

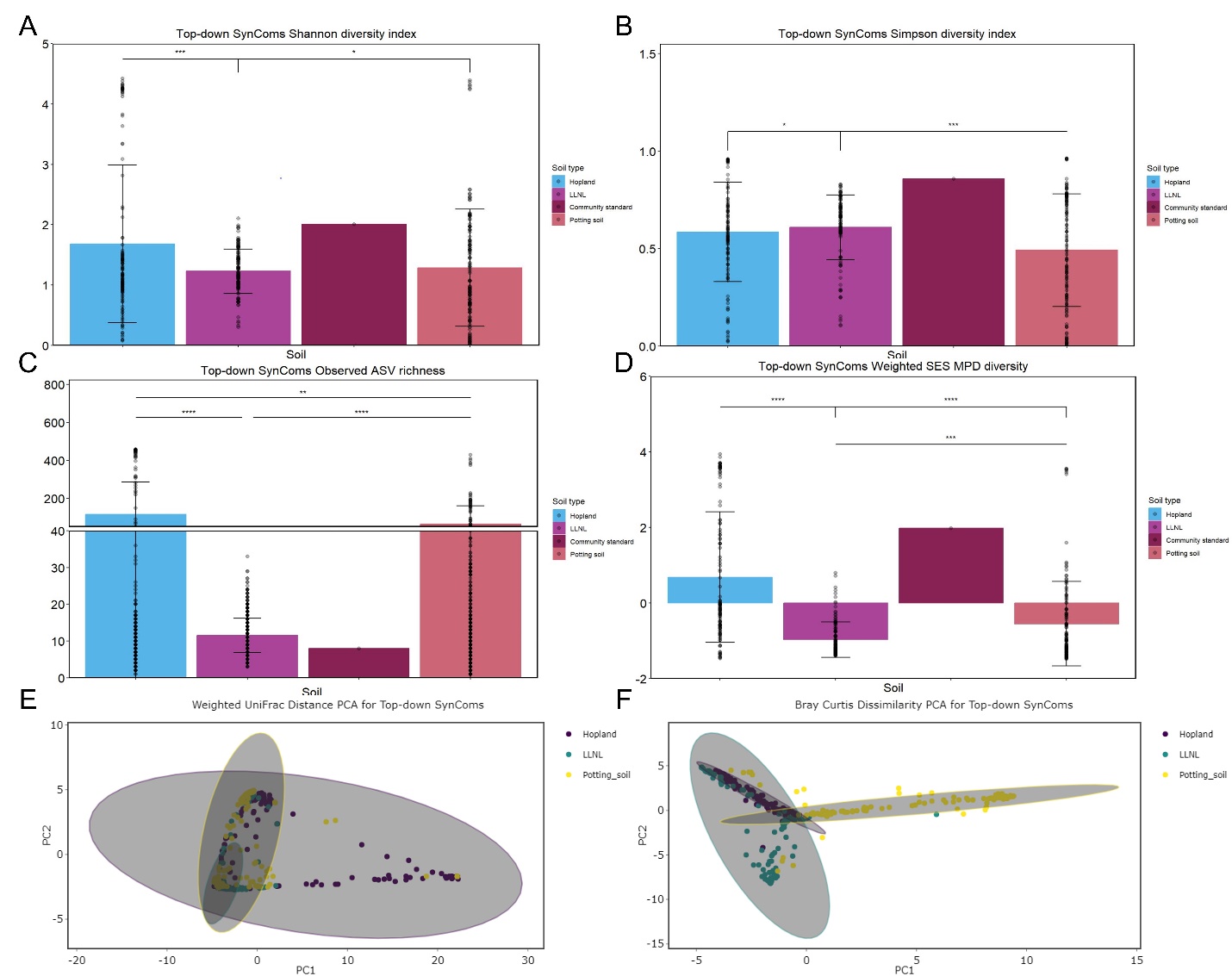

**Figure 3**. **Soil level breakdown of top-down SynComs reveal trends in diversity, richness and beta diversity grouping**

The base or control top-down SynComs were assessed at the soil level using Shannon diversity index (A), Simpson diversity index (B) and standardized effect size (SES) of mean pairwise distance (MPD) phylogenetic diversity (D) for alpha diversity. Observed amplicon sequencing variant (ASV) richness (C) was used for richness estimation and Weighted UniFrac distance (E) and Bray Curtis Dissimilarity (F) are shown for beta diversity as PCA plots with the ellipses representing 95% confidence intervals. Data for alpha diversity is mean ± SD with each SynCom represented as a point which is on average 133 individual replicates as on average 66 SynComs in every soil and two separate 96-well plates were sequenced for each soil condition with the same for PCA plots; *p<0.05, **p<0.01, ***p<0.001, ****p<0.0001.

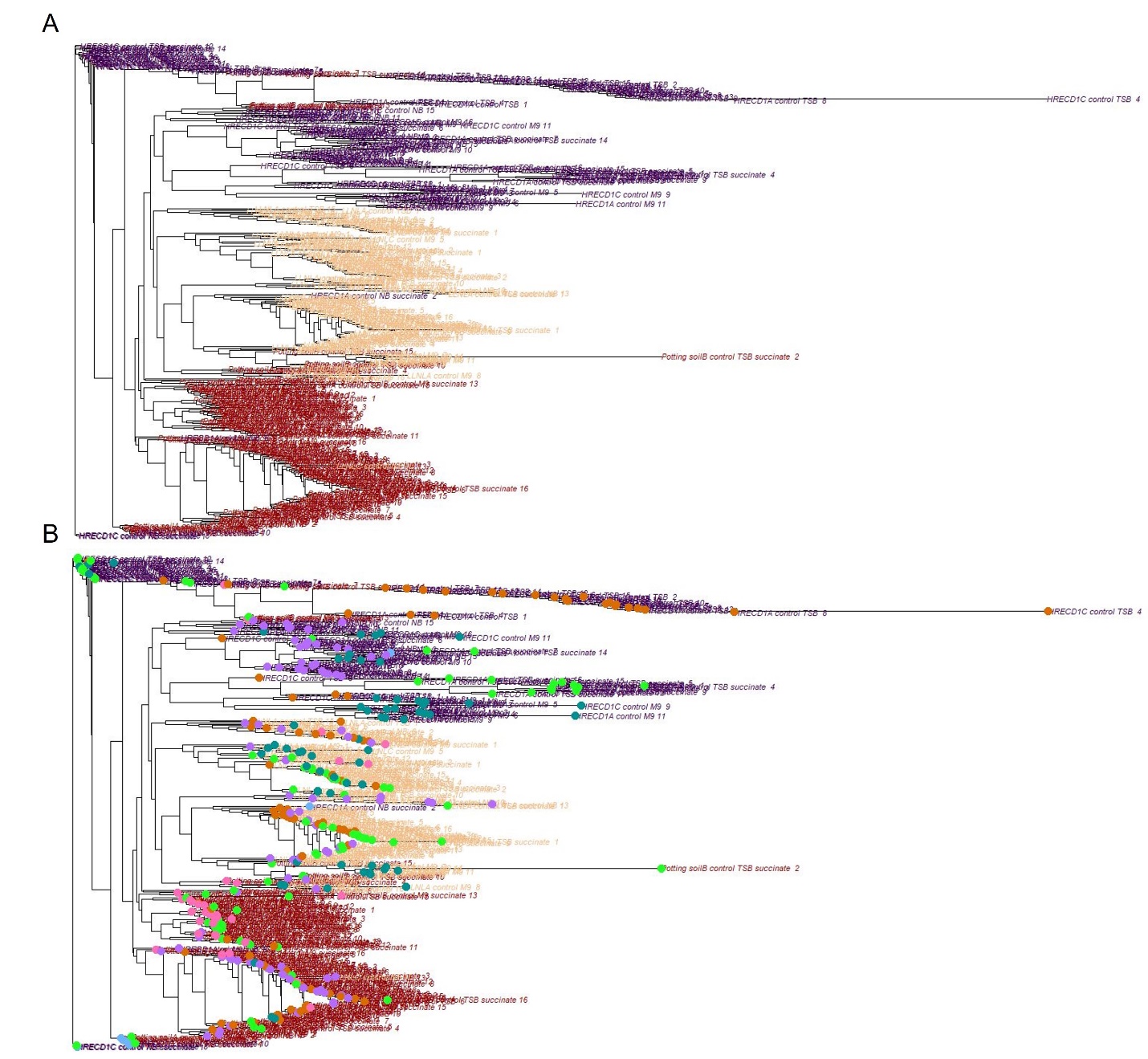

**Figure 4**. **Phylogenetic trees at the community level emphasize trends of grouping based on soil type with more distribution regarding media type**

The base or control top-down SynComs were constructed into rooted phylogenetic trees at the community level. The designation of control is applied to these SynComs as these are the base SynComs that the two *B. subtilis* strains were inoculated into generating the other SynComs. The first phylogenetic tree was colored based on soil type (A) with Hopland (HRECD1) colored in purple, LLNL colored in beige and Potting soil colored in maroon. A second tree (B) was constructed where the soil types are still colored the same as the first then the media types were overlayed onto the tree using colored dots that correlate to the media type colors in the other graphs. The media type colors include turquoise (M9), pink (M9 succinate), purple (NB), light blue (NB succinate), orange (TSB) and light green (TSB succinate).

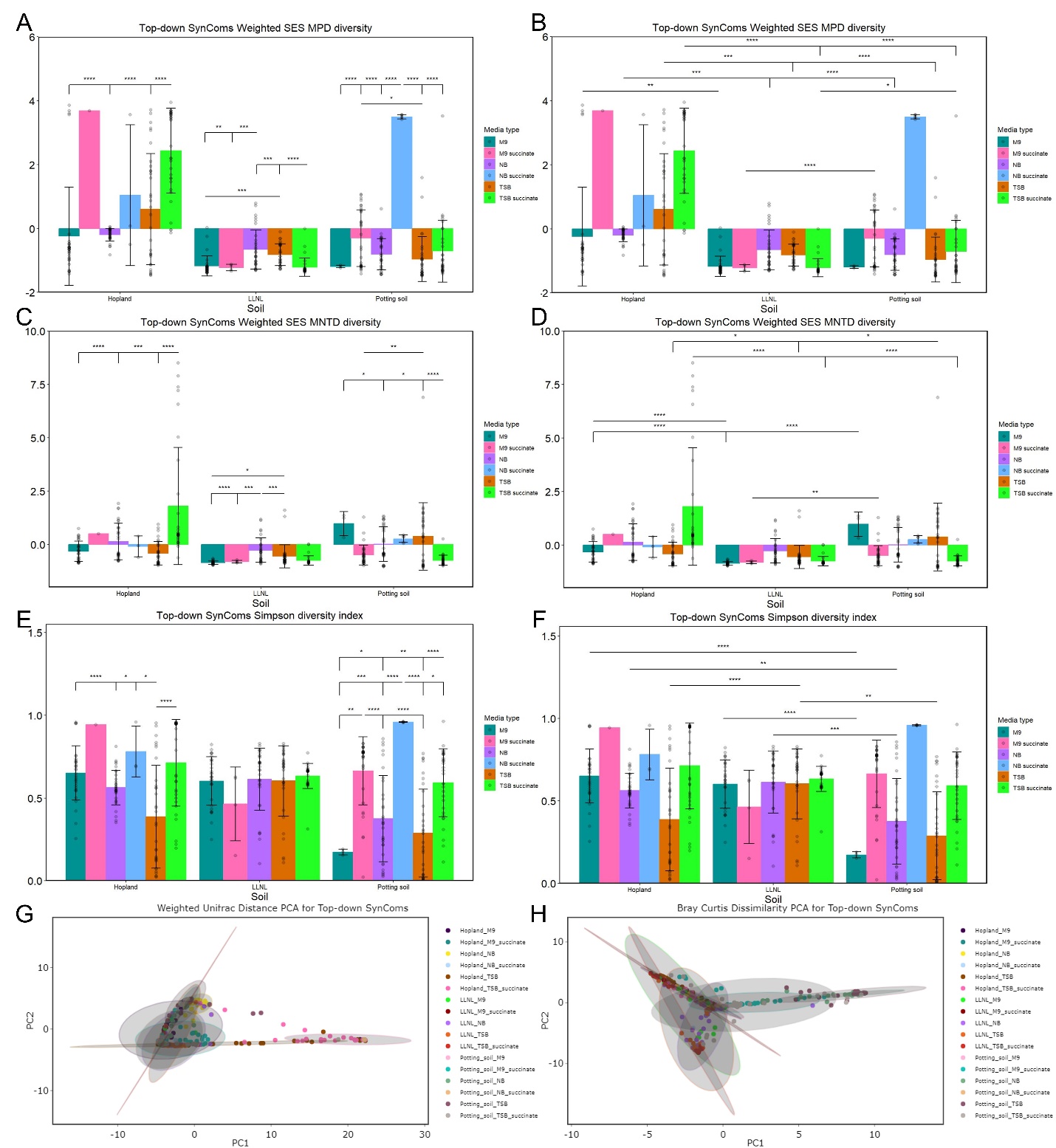

**Figure 5**. **Further analyses and visualizations reinforce observed trends in top-down SynComs from other metrics**

The base or control top-down SynComs were further assessed at the soil and media type level using standardized effect size (SES) of mean pairwise distance (MPD) and SES mean nearest taxon distance (MNTD) phylogenetic diversity (A, B, C, D) and Simpson diversity index (E, F) for alpha diversity. Intra-soil statistics shown on the left (A, C, E) and inter-soil statistics shown on the right (B, D, F). Weighted UniFrac distance (G) and Bray Curtis Dissimilarity (H) are shown for beta diversity as PCA plots with the ellipses representing 95% confidence intervals. Data for alpha diversity is mean ± SD with each SynCom represented as a point which is up to 32 individual replicates as there were 16 SynComs in each media type of every soil and two separate 96-well plates were sequenced for every condition with the same for PCA plots; *p<0.05, **p<0.01, ***p<0.001, ****p<0.0001.

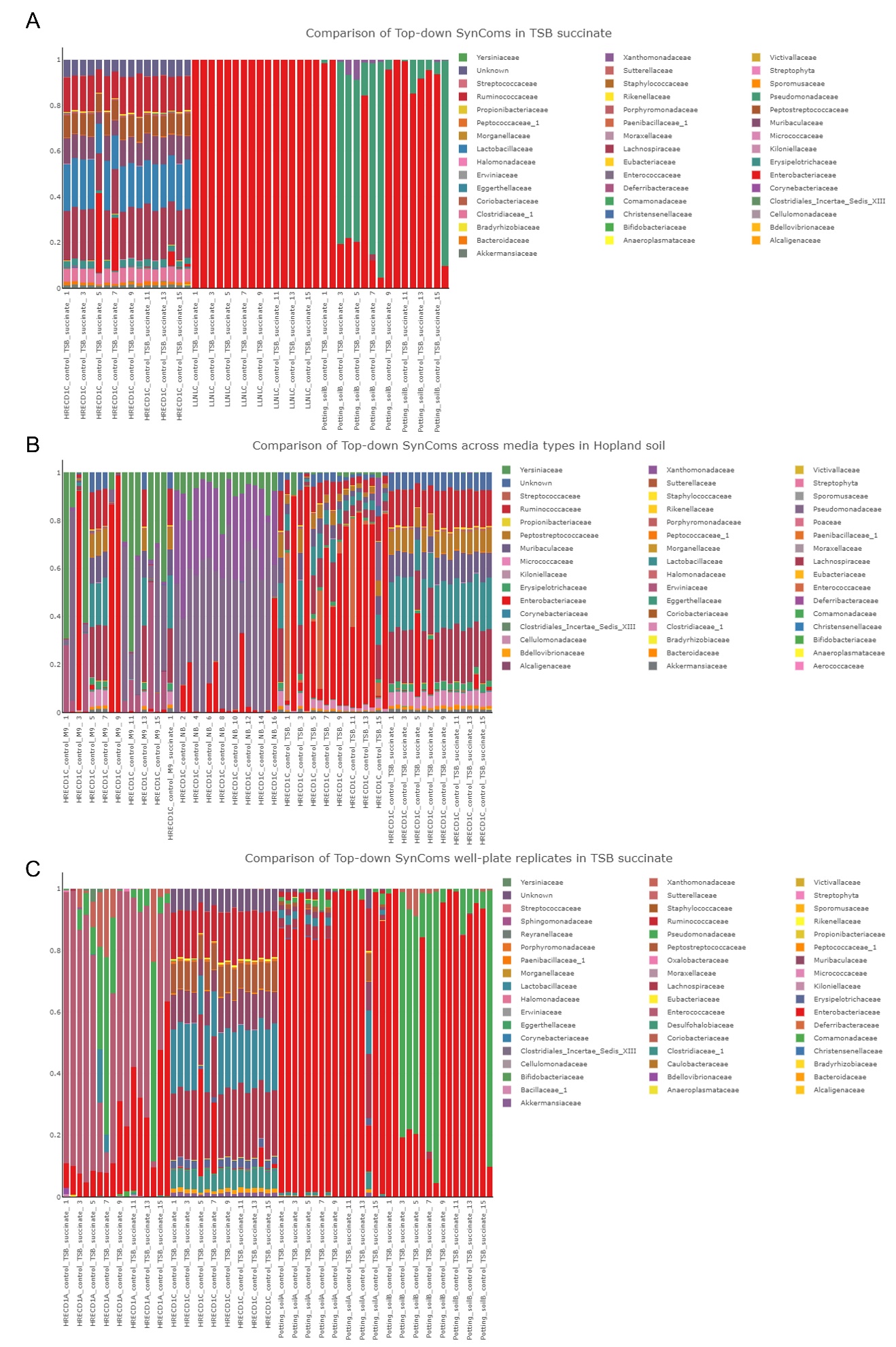

**Figure 6**. **Representative composition profiles explain complexity observed in alpha and beta diversity quantifications**

Three representative composition profiles at the family level were assembled to display various trends observed in the SynComs and contextualize the results seen in the other metrics. The designation of control is applied to these SynComs as these are the base SynComs that the two *B. subtilis* strains were inoculated into generating the other SynComs. The first compositional profile was used to compare diversity, richness and community structure of Hopland (HRECD1), LLNL and Potting soil in the TSB succinate media type (A). The second compositional profile was used to compare all the media types from the SynComs sequenced in Hopland as a representation of the complexity established in different media types from the same soil (B). The third compositional profile was used to demonstrate the consistency that can be established between what might be considered “technical” replicates (same soil and media type from the same well plate) and how different “biological” replicates (same soil and media type from different well plates) can be from different 5 grams of the original soil (C).

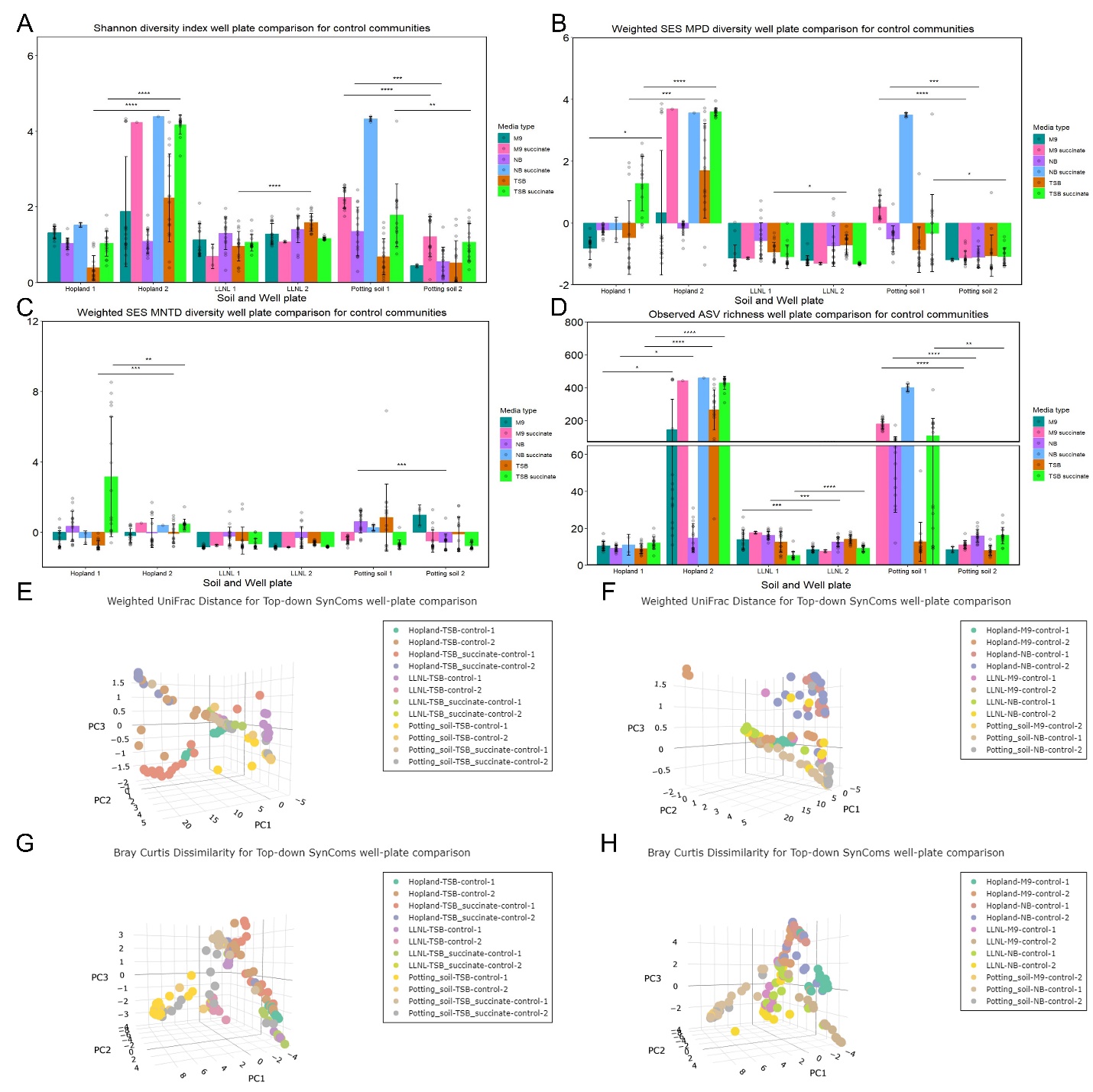

**Figure 7**. **SynComs from separate well-plates that were sourced and propagated from the same soil and media type showed consistency and variability based on soil and media type**

The base or control top-down SynComs originating from each 96-well plate (“biological replicates”) are visualized using several metrics for alpha and beta diversity. Shannon diversity index (A), standardized effect size (SES) of mean pairwise distance (MPD) and SES mean nearest taxon distance (MNTD) phylogenetic diversities (B, C) are shown for alpha diversity with Observed amplicon sequencing variant (ASV) richness (D) used for estimating richness. Intra-soil statistics are shown due to comparing between the “biological replicates”. For beta diversity, Weighted UniFrac distance (E, F) and Bray Curtis Dissimilarity (G, H) are visualized as PCA plots with some media types split between left and right side for easier visualization. The designation of control is applied to these SynComs as these are the base SynComs that the two *B. subtilis* strains were inoculated into generating the other SynComs. Data for alpha diversity is mean ± SD with each SynCom represented as a point which could be up to 16 individual replicates as the 16 SynComs in each media type of every soil in the two separate 96-well plates are being compared with the same for PCA plots; *p<0.05, **p<0.01, ***p<0.001, ****p<0.0001.

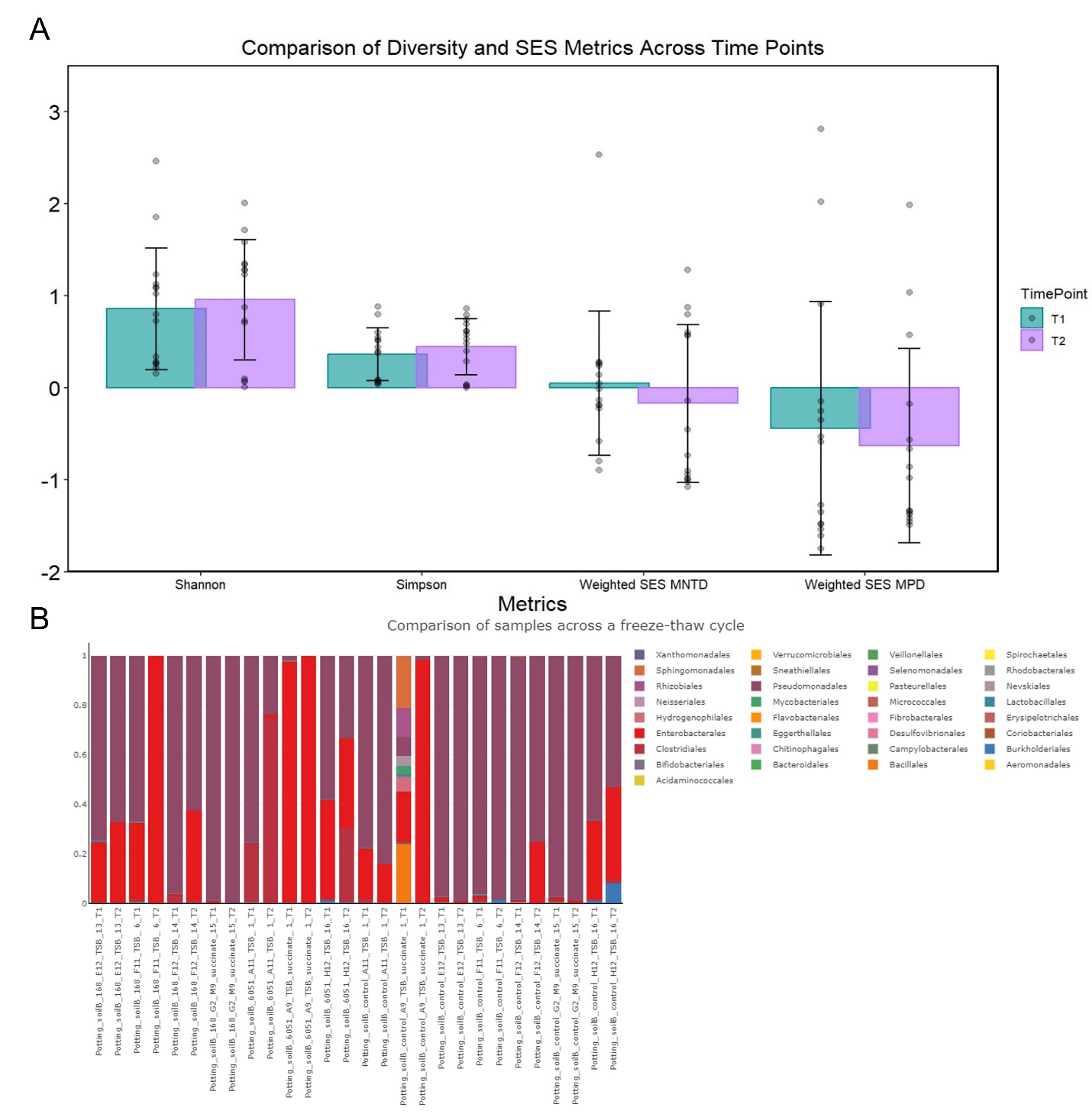

**Figure 8**. **SynComs overall maintained stability across a storage and reanimation process**

Representative base or control top-down SynComs that were sequenced across one deep freeze storage then regrowth are visualized using several metrics for alpha diversity and a composition profile to determine stability through this process. Shannon diversity index, Simpson diversity index, standardized effect size (SES) of mean nearest taxon distance (MNTD) and SES mean pairwise distance (MPD) phylogenetic diversities are shown for alpha diversity (A). Composition profile at the order level is shown to compare across this storage and reanimation process (B). Before and after this process is referred to as T1 and T2 respectively with T2 being the same time as all other sequencing. The designation of control is applied to these SynComs as these are the base SynComs that the two *B. subtilis* strains were inoculated into generating the other SynComs. Data for alpha diversity is mean ± SD with each SynCom represented as a point which is 15 individual replicates at each time point.

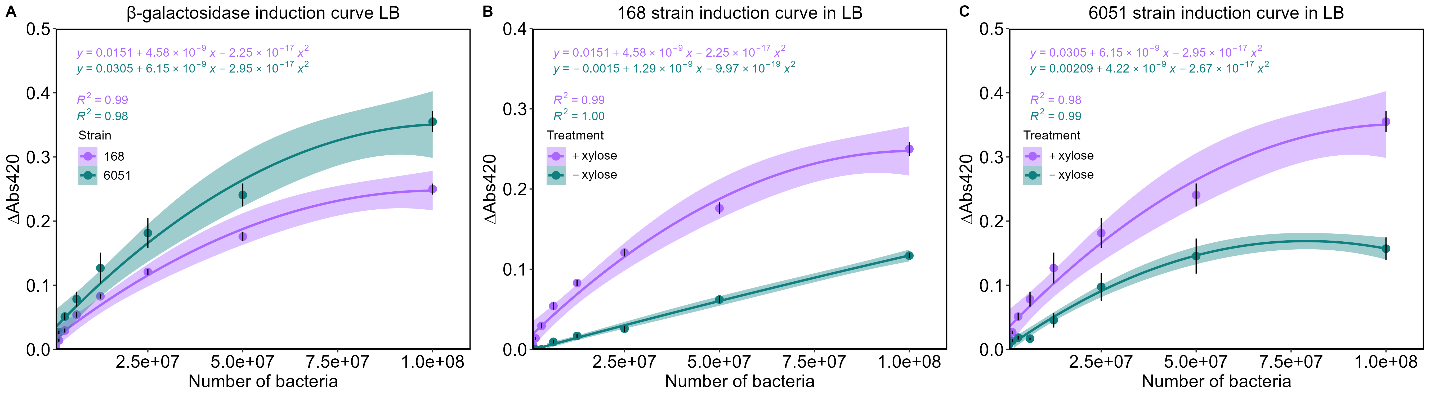

**Figure 9**. ***B. subtilis* strains express β-galactosidase when controlled by xylose system**

*B. subtilis* 168 and 6051a (6051) strains expressing β-galactosidase under control of the xylose inducible system were serially diluted and induced or not induced with xylose in the conditions that would be used for screening in the SynComs to test change in absorbance after adding ONPG in LB. Best fit polynomial equations were generated in R to connect change in absorbance to abundance of each *B. subtilis* strain. The 168 and 6051a strains are compared after induction (A), 168 strain with or without xylose (B) and 6051a strain with or without xylose (C). Data is mean ± SEM of 12 replicates (n = 3 biological, n = 4 technical).

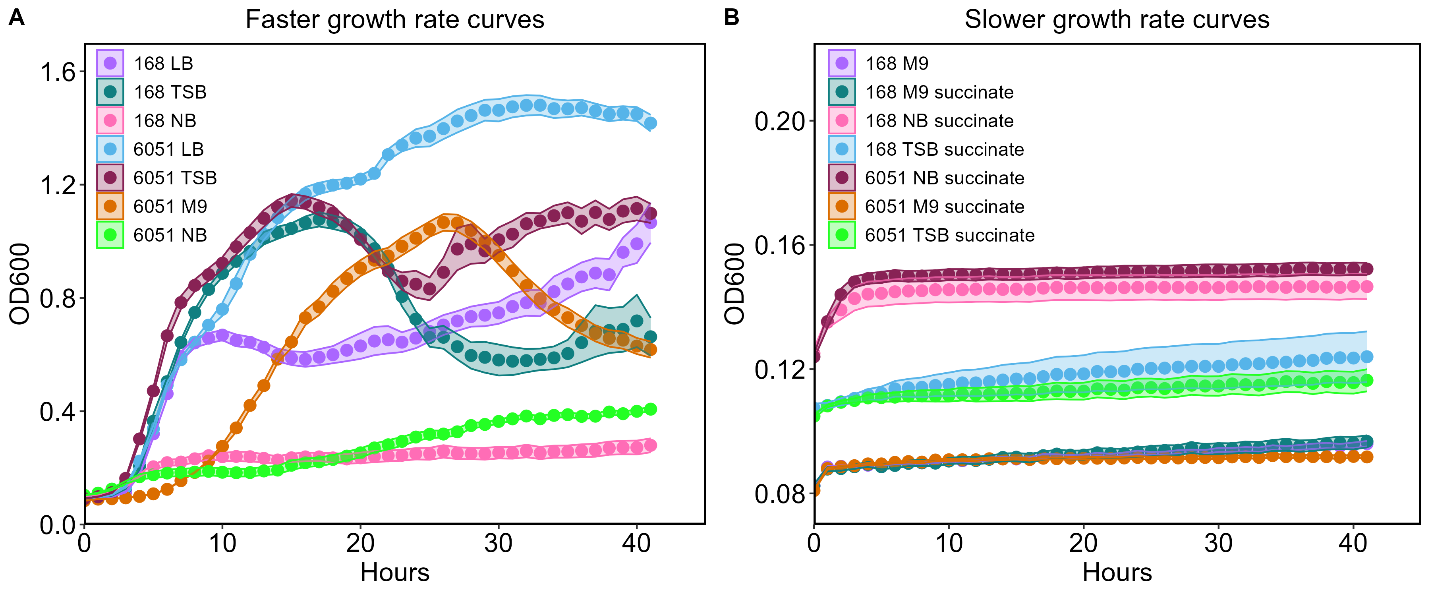

**Figure 10**. **Growth curves for 168 and 6051a strains in all media conditions**

*B. subtilis* 168 and 6051a (6051) strains were grown in all media types associated with SynCom growth and LB was used as a control for standard culture conditions. All cultures were normalized to OD_600_ = 1 in PBS then a 1:100 was performed into every media type. Culture conditions where the *B. subtilis* strains had faster growth conditions are shown together (A) and those conditions where the strains exhibited less growth are shown together (B). Data is mean ± SEM of 12 replicates (n = 3 biological, n = 4 technical).

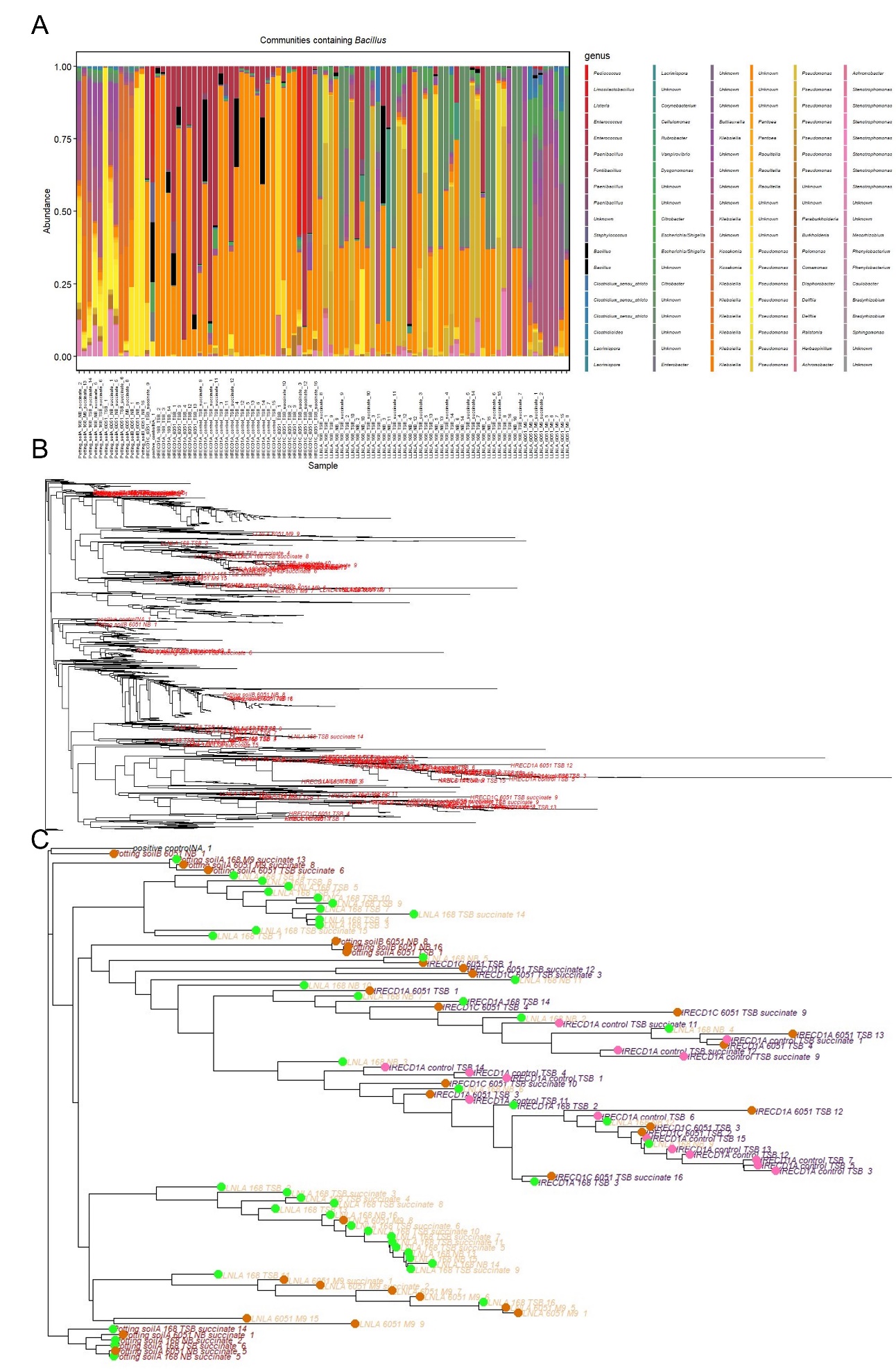

**Figure 11**. ***B. subtilis* 168 and 6051a strains persisted in a variety of SynComs from multiple soil and media types**

Composition profile at the genus level highlighting only SynComs with *Bacillus* ASVs (black) was assembled after all base SynComs and SynComs inoculated with the *B. subtilis* strains were analyzed post 16S rRNA sequencing with Potting soil on the left, Hopland (HRECD1, only condition with *Bacillus* in some base SynComs) in the middle and LLNL on the right (A). All of the SynComs (control and inoculated conditions) were constructed into a rooted phylogenetic tree at the community level with SynComs containing *Bacillus* highlighted in red to symbolize the phylogenetic diversity of *Bacillus* containing SynComs (B). A second tree at the community level (C) was constructed of just the *Bacillus* containing SynComs where the soil types are Hopland (HRECD1) colored in purple, LLNL colored in beige and Potting soil colored in maroon then the inoculation type was overlayed onto the tree using colored dots: control or base SynCom (pink), 168 strain (light green) and 6051a (6051) strain (orange).

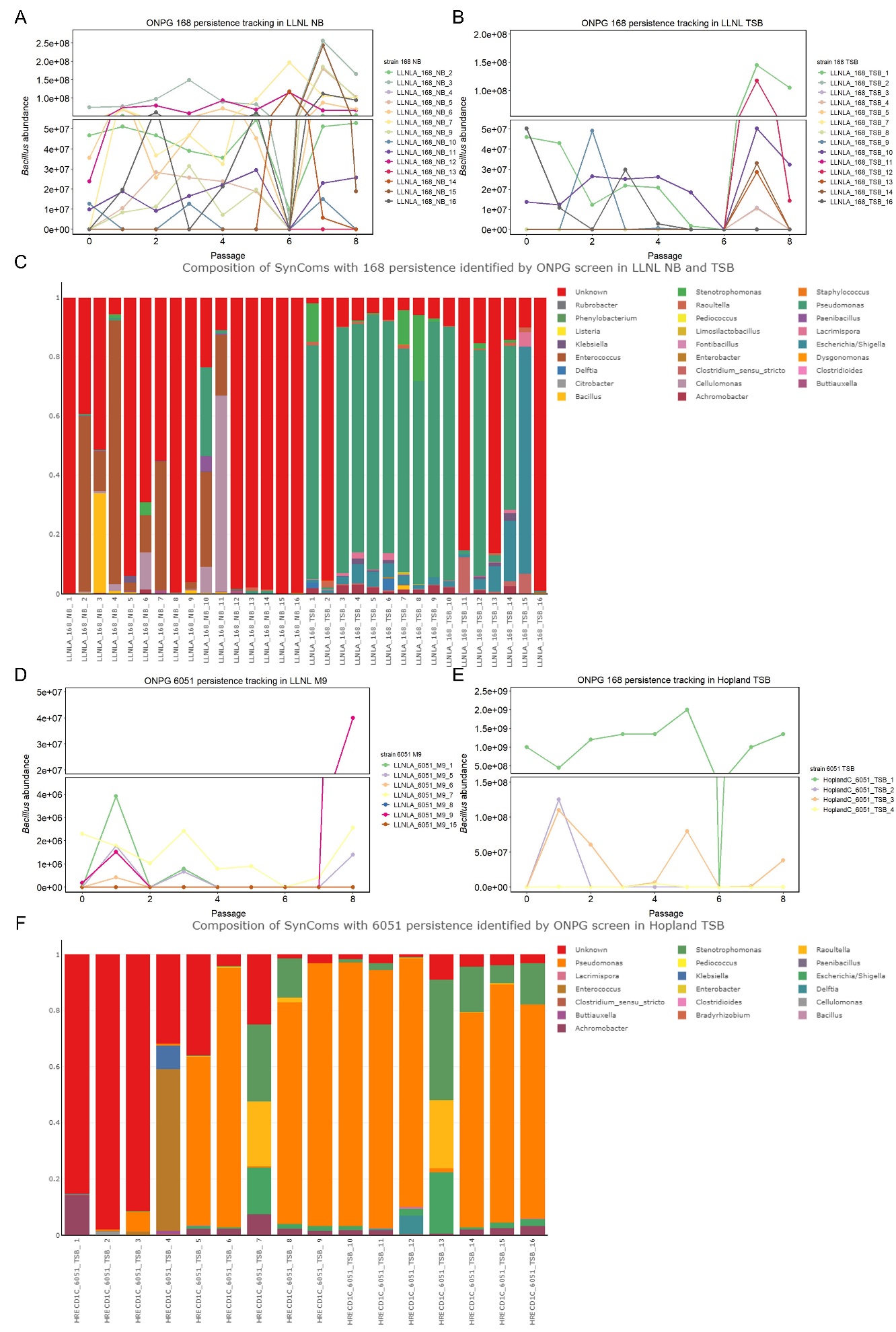

**Figure 12**. ***Bacillus* abundance tracking curves provided insight into relative abundance found in the composition profiles especially for 168 strain in LLNL NB**

Abundance tracking curves were constructed by converting absorbance of SynComs that the 168 and 6051a (6051) strain were inoculated into after subtracting background from the original base SynComs without inoculation into abundance by using the calibration curves in each of the SynCom media types. The abundance tracking curves are plotted based on soil, media and inoculation type with 168 strain conditions (A, B) and 6051a (D, E) separated on top and bottom. In conditions where absorbance was outside of the calibration range, the abundance was estimated based on the calibration curves. Composition profiles at the genus level for soil, media and inoculation conditions in the abundance tracking curves are shown with 168 strain in LLNL conditions on in the middle (C) and the 6051a strain in Hopland (HRECD1) conditions on the bottom (F).

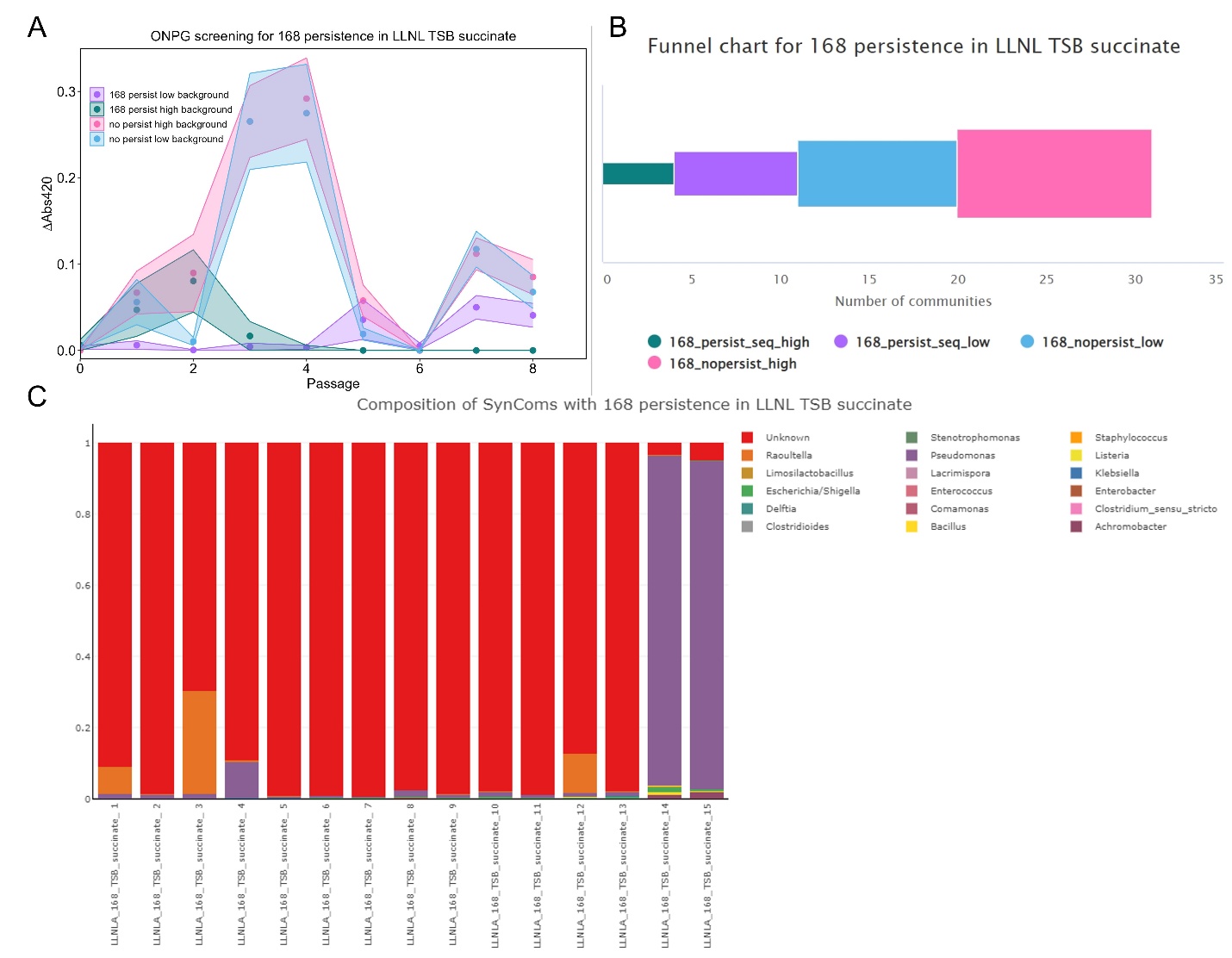

**Figure 13**. **ONPG screen did not distinguish persistence for 168 strain in LLNL TSB succinate**

Temporal tracking curves were constructed by plotting change in absorbance of SynComs that the 168 strain was inoculated into after subtracting background from the original base SynComs without inoculation. The SynComs were classified into four groups after persistence was revealed or not revealed by sequencing (persist or no persist). High or low background was determined by whether the original base SynCom was quantified to have absorbance values greater or less than the maximum value for each strain in each media type in the calibration curves. Tracking curve (A) and funnel chart (B) are shown for the specific soil and media type with funnel chart designed to represent the number of SynComs that were grouped into each subcategory based on the downselection process of the ONPG screen and sequencing. Colors for categories are the same between both types of plots (sequencing indicated—persist_seq, not indicated—nopersist; high background—_high, low background—_low). Points in the tracking curve is mean ± SEM of the number of replicates shown in the funnel chart based on how the SynComs were grouped. Composition profile (C) at the genus level where persistence was shown by sequencing is plotted.

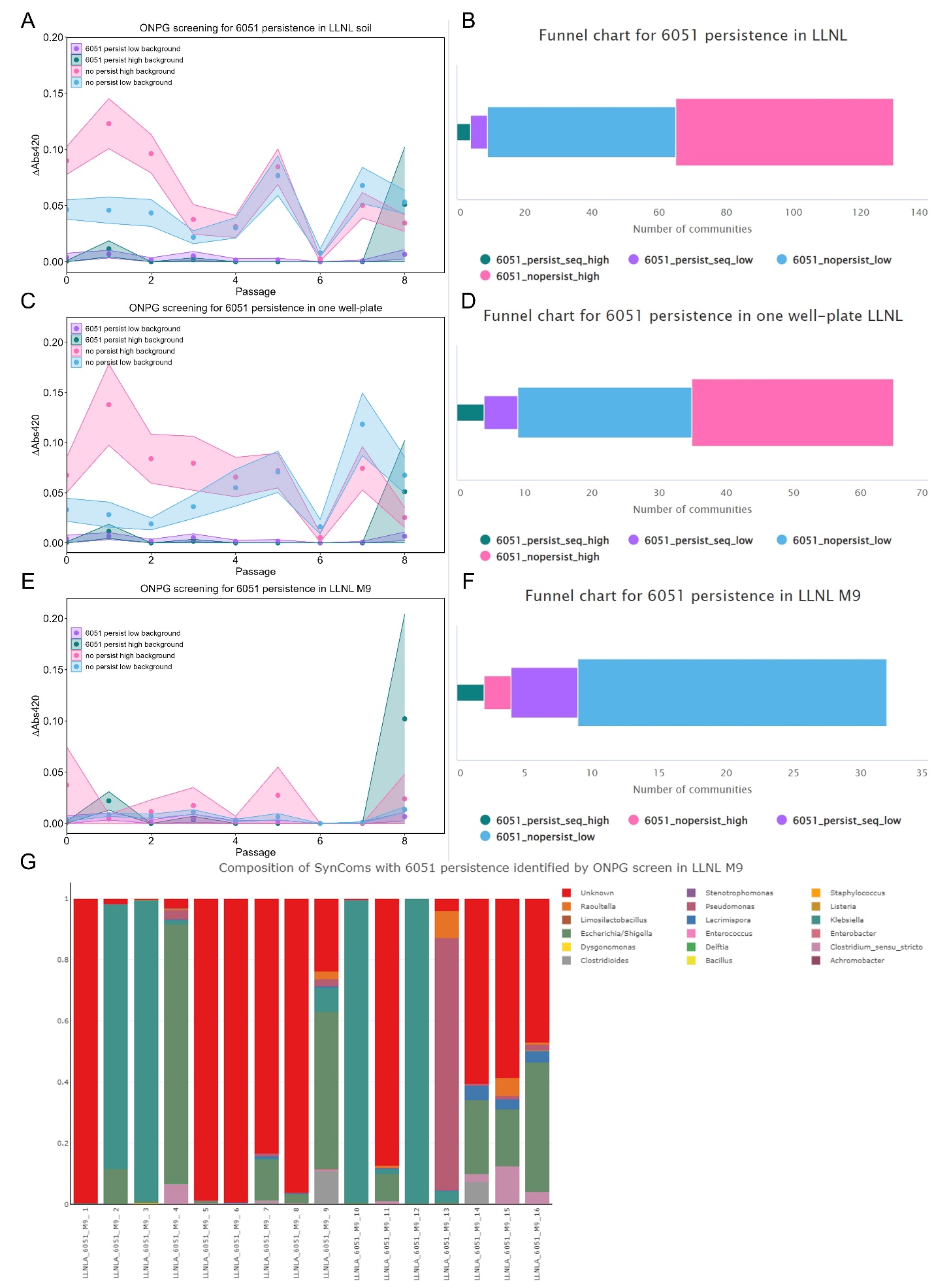

**Figure 14**. **ONPG screen identified possible SynComs in LLNL conditions where the 6051a strain expression of β-gal was used for downselection**

Temporal tracking curves were constructed by plotting change in absorbance of SynComs that the 6051a (6051) strain was inoculated into after subtracting background from the original base SynComs without inoculation. The SynComs were classified into four groups after persistence was revealed or not revealed by sequencing (persist or no persist). High or low background was determined by whether the original base SynCom was quantified to have absorbance values greater or less than the maximum value for each strain in each media type in the calibration curves. The tracking curves are plotted based on soil (A), then subdivided to just one well plate (c) or subdivided into media type (E). Funnel charts (B, D, F) which are paired with the adjoining tracking curves are shown to represent the number of SynComs that were grouped into each subcategory based on the downselection process of the ONPG screen and sequencing. Colors for categories are the same between both types of plots (sequencing indicated—persist_seq, not indicated—nopersist; high background—_high, low background—_low). Points in the tracking curve is mean ± SEM of the number of replicates shown in the funnel chart based on how the SynComs were grouped. Composition profile (C) at the genus level where persistence was shown by sequencing is plotted.

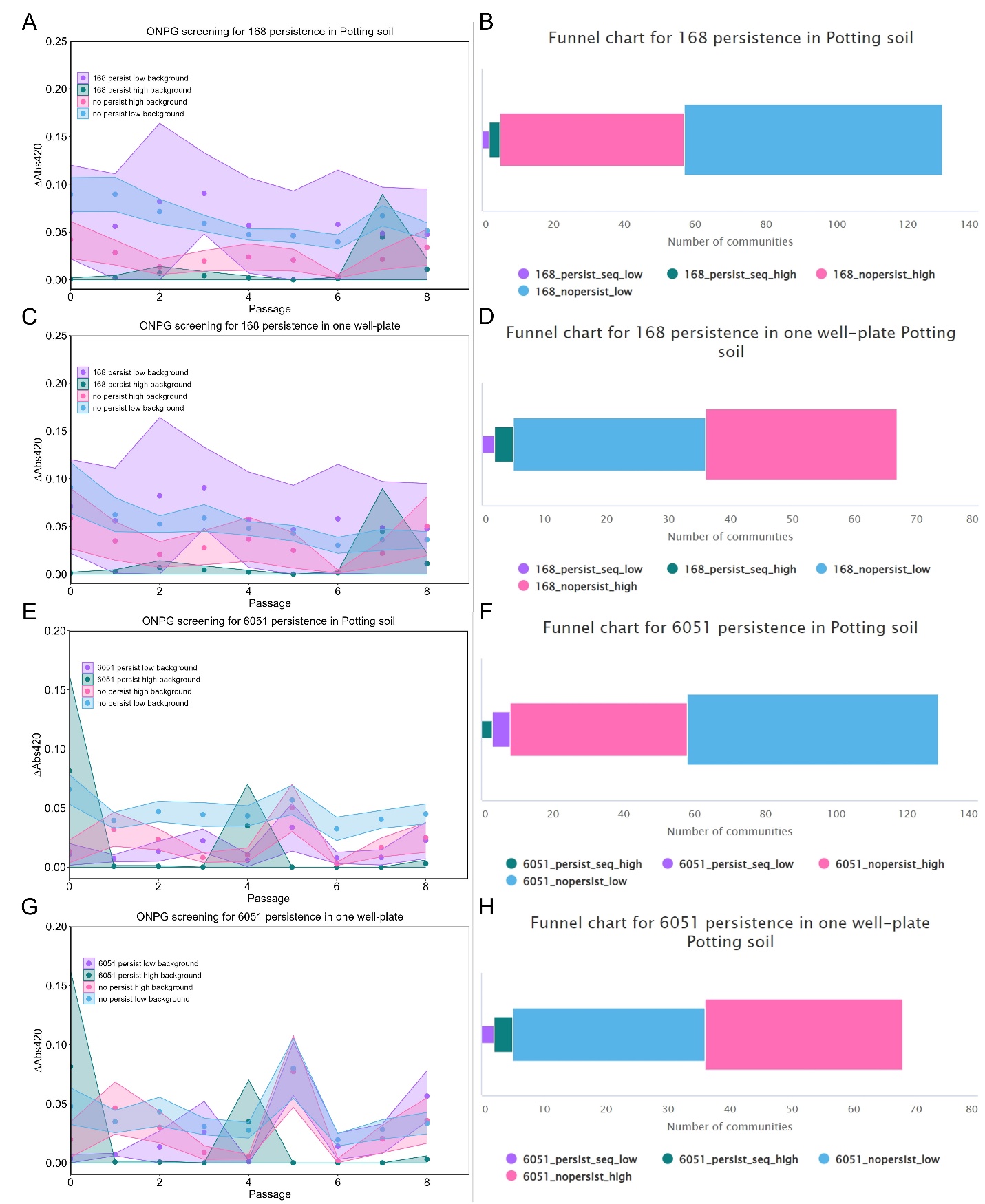

**Figure 15**. **High variance in all conditions was observed during the ONPG screen of Potting soil for both strains**

Temporal tracking curves were constructed by plotting change in absorbance of SynComs that the 168 and 6051a (6051) strains were inoculated into after subtracting background from the original base SynComs without inoculation. The SynComs were classified into four groups after persistence was revealed or not revealed by sequencing (persist or no persist). High or low background was determined by whether the original base SynCom was quantified to have absorbance values greater or less than the maximum value for each strain in each media type in the calibration curves. The tracking curves are plotted based on soil (A, E) then subdivided to just one well plate (C, G). Funnel charts (B, D, F, H) which are paired with the adjoining tracking curves are shown to represent the number of SynComs that were grouped into each subcategory based on the downselection process of the ONPG screen and sequencing. Colors for categories are the same between both types of plots (sequencing indicated—persist_seq, not indicated—nopersist; high background—_high, low background—_low). Points in the tracking curve is mean ± SEM of the number of replicates shown in the funnel chart based on how the SynComs were grouped.

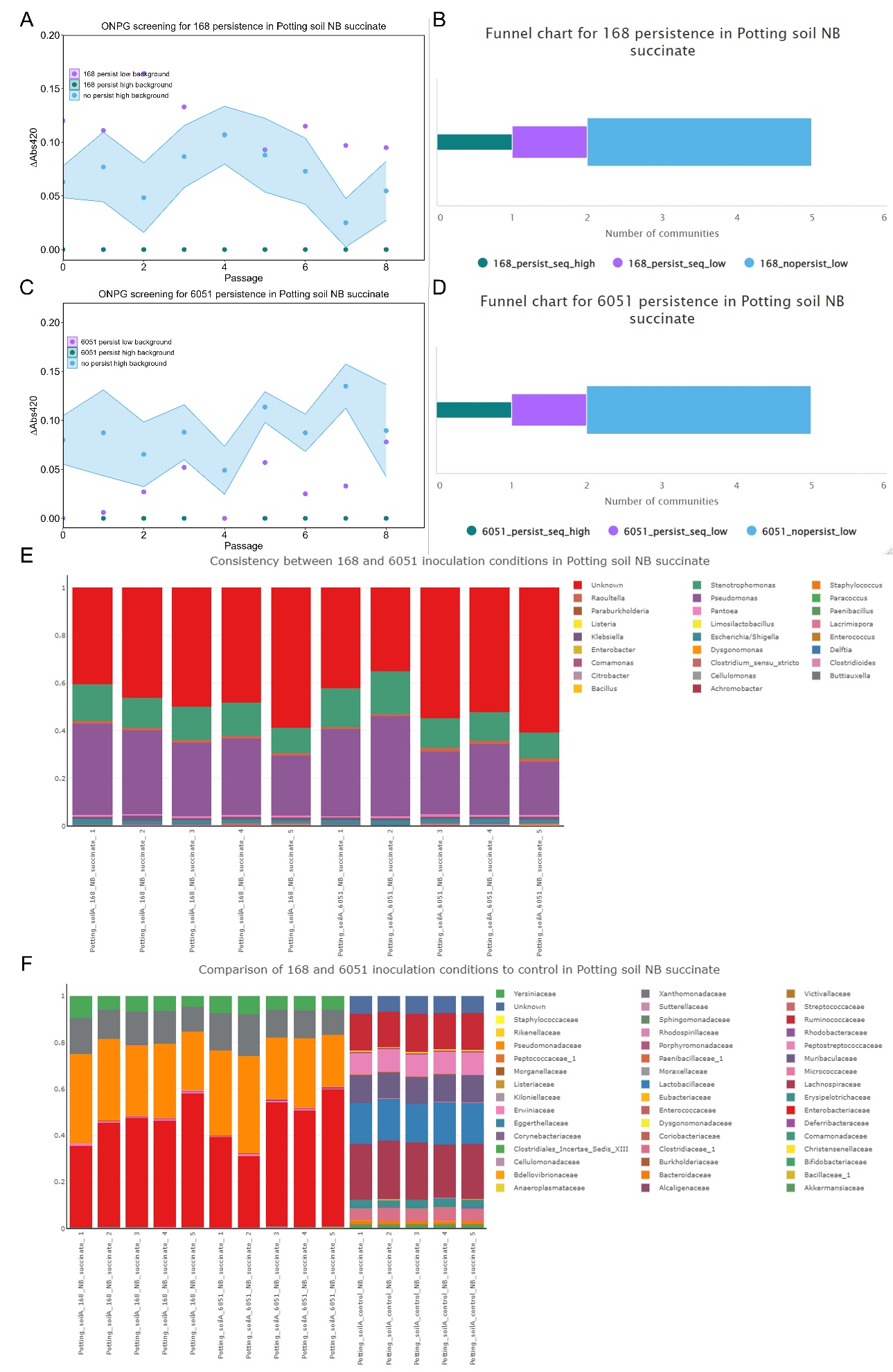

**Figure 16**. **ONPG screen and sequencing identified persistence in the same soil and media condition for both strains**

Temporal tracking curves were constructed by plotting change in absorbance of SynComs that the 168 and 6051a (6051) strains were inoculated into after subtracting background from the original base SynComs without inoculation. The SynComs were classified into four groups after persistence was revealed or not revealed by sequencing (persist or no persist). High or low background was determined by whether the original base SynCom was quantified to have absorbance values greater or less than the maximum value for each strain in each media type in the calibration curves. The tracking curves are plotted based on specific soil and media type (A, C). Funnel charts (B, D) which are paired with the adjoining tracking curves are shown to represent the number of SynComs that were grouped into each subcategory based on the downselection process of the ONPG screen and sequencing. Colors for categories are the same between both types of plots (sequencing indicated—persist_seq, not indicated—nopersist; high background—_high, low background—_low). Points in the tracking curve is mean ± SEM of the number of replicates shown in the funnel chart based on how the SynComs were grouped. Composition profiles at the genus level comparing SynComs from the 168 and 6051 persistence (E) and at the family level comparing SynComs from 168, 6051 and the base SynComs (F) are plotted.

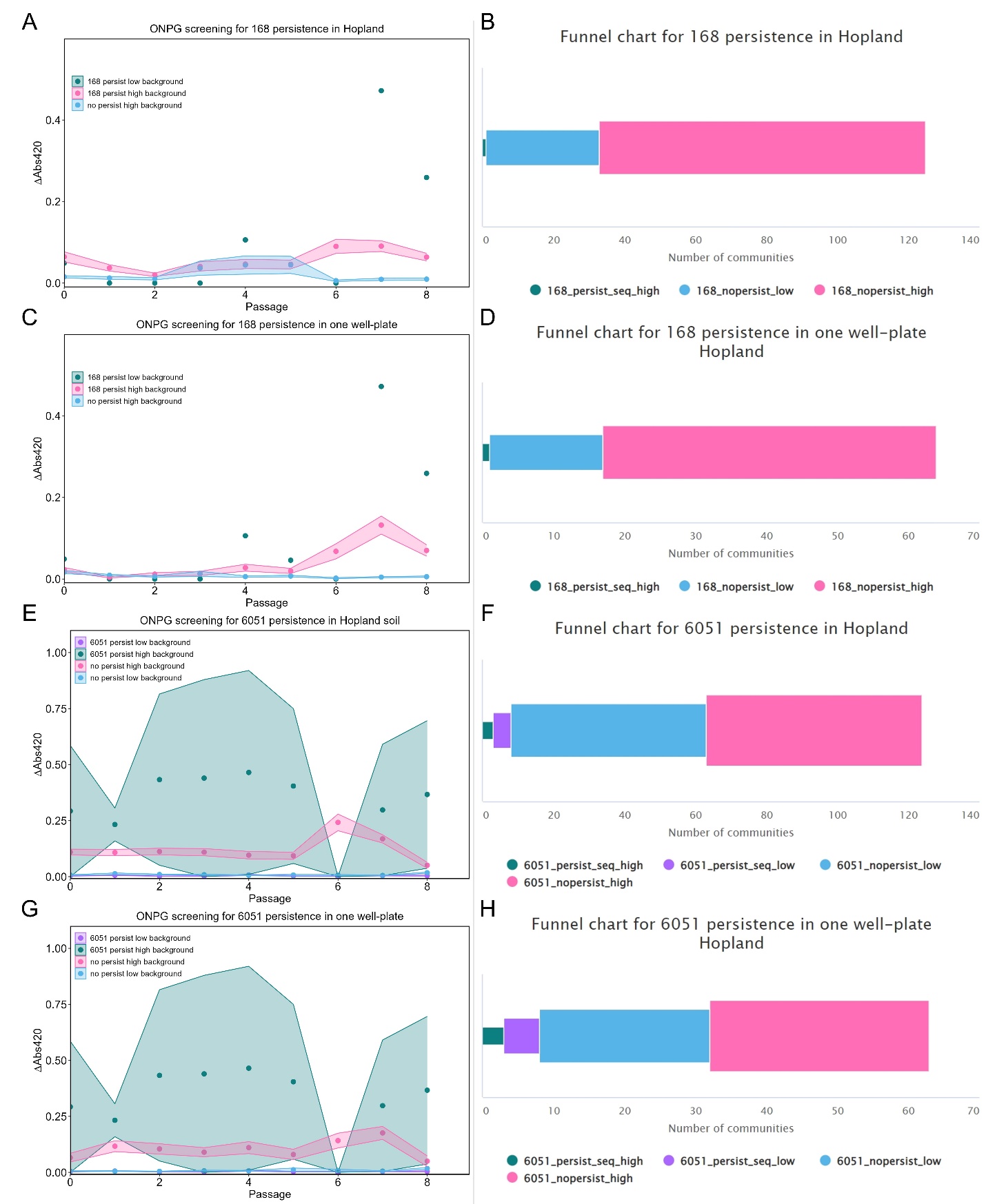

**Figure 17**. **ONPG screen identified further SynComs in Hopland conditions where the 168 and 6051a strain expression of β-gal supported downselection for sequencing**

Temporal tracking curves were constructed by plotting change in absorbance of SynComs that the 168 and 6051a (6051) strains were inoculated into after subtracting background from the original base SynComs without inoculation. The SynComs were classified into four groups after persistence was revealed or not revealed by sequencing (persist or no persist). High or low background was determined by whether the original base SynCom was quantified to have absorbance values greater or less than the maximum value for each strain in each media type in the calibration curves. The tracking curves are plotted based on soil (A, E) then subdivided to just one well plate (C, G). Funnel charts (B, D, F, H) which are paired with the adjoining tracking curves are shown to represent the number of SynComs that were grouped into each subcategory based on the downselection process of the ONPG screen and sequencing. Colors for categories are the same between both types of plots (sequencing indicated—persist_seq, not indicated—nopersist; high background—_high, low background—_low). Points in the tracking curve is mean ± SEM of the number of replicates shown in the funnel chart based on how the SynComs were grouped.

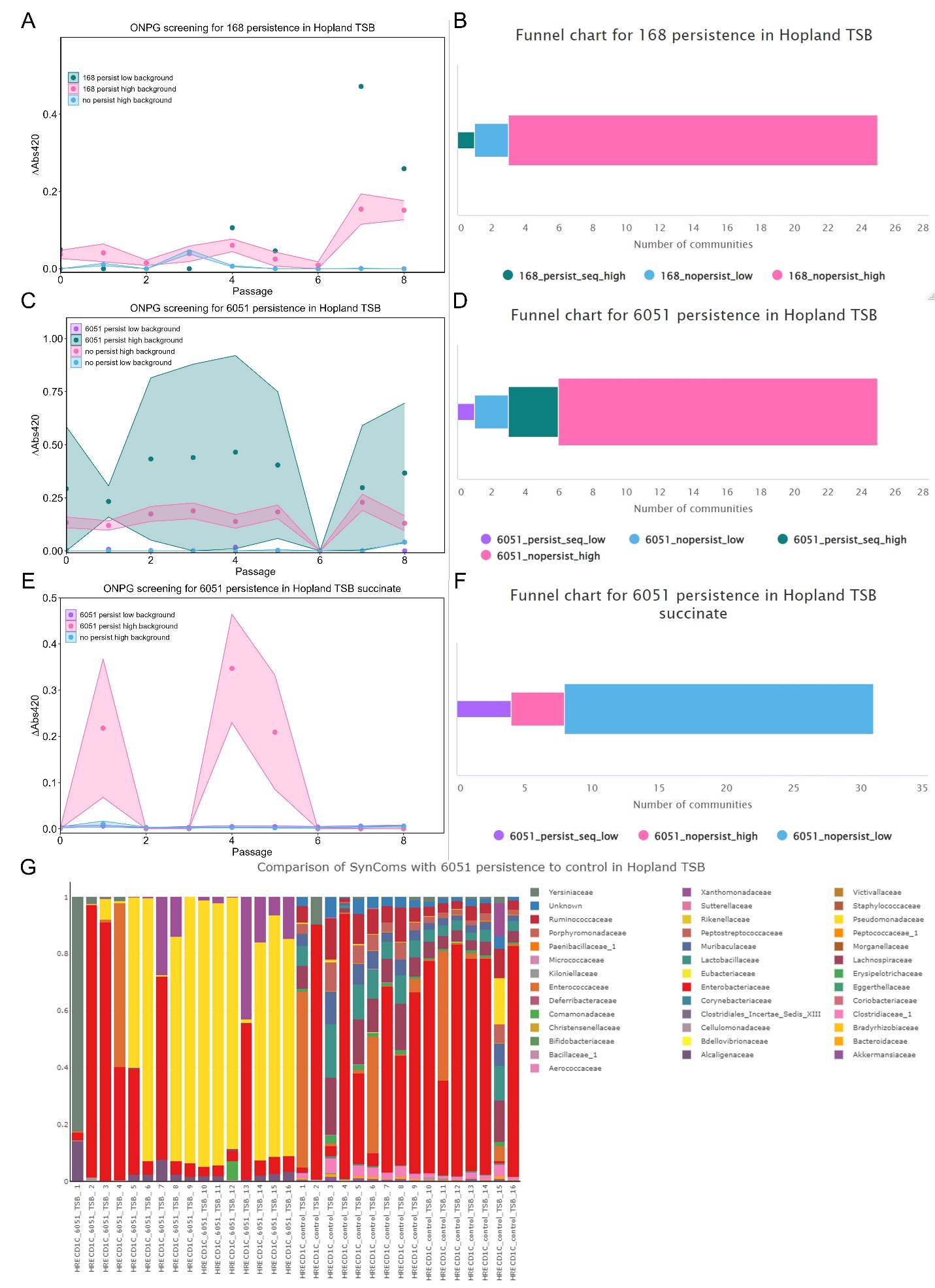

**Figure 18**. **Specific media conditions correlated to the SynComs that separated in the ONPG screen in Hopland**

Temporal tracking curves were constructed by plotting change in absorbance of SynComs that the 168 and 6051a (6051) strains were inoculated into after subtracting background from the original base SynComs without inoculation. The SynComs were classified into four groups after persistence was revealed or not revealed by sequencing (persist or no persist). High or low background was determined by whether the original base SynCom was quantified to have absorbance values greater or less than the maximum value for each strain in each media type in the calibration curves. The tracking curves are plotted based on media type for both 168 (A) and 6051a (C, E). Funnel charts (B, D, F) which are paired with the adjoining tracking curves are shown to represent the number of SynComs that were grouped into each subcategory based on the downselection process of the ONPG screen and sequencing. Colors for categories are the same between both types of plots (sequencing indicated—persist_seq, not indicated—nopersist; high background—_high, low background—_low). Points in the tracking curve is mean ± SEM of the number of replicates shown in the funnel chart based on how the SynComs were grouped. Composition profile (g) at the family level where 6051a persistence was shown by sequencing is plotted.

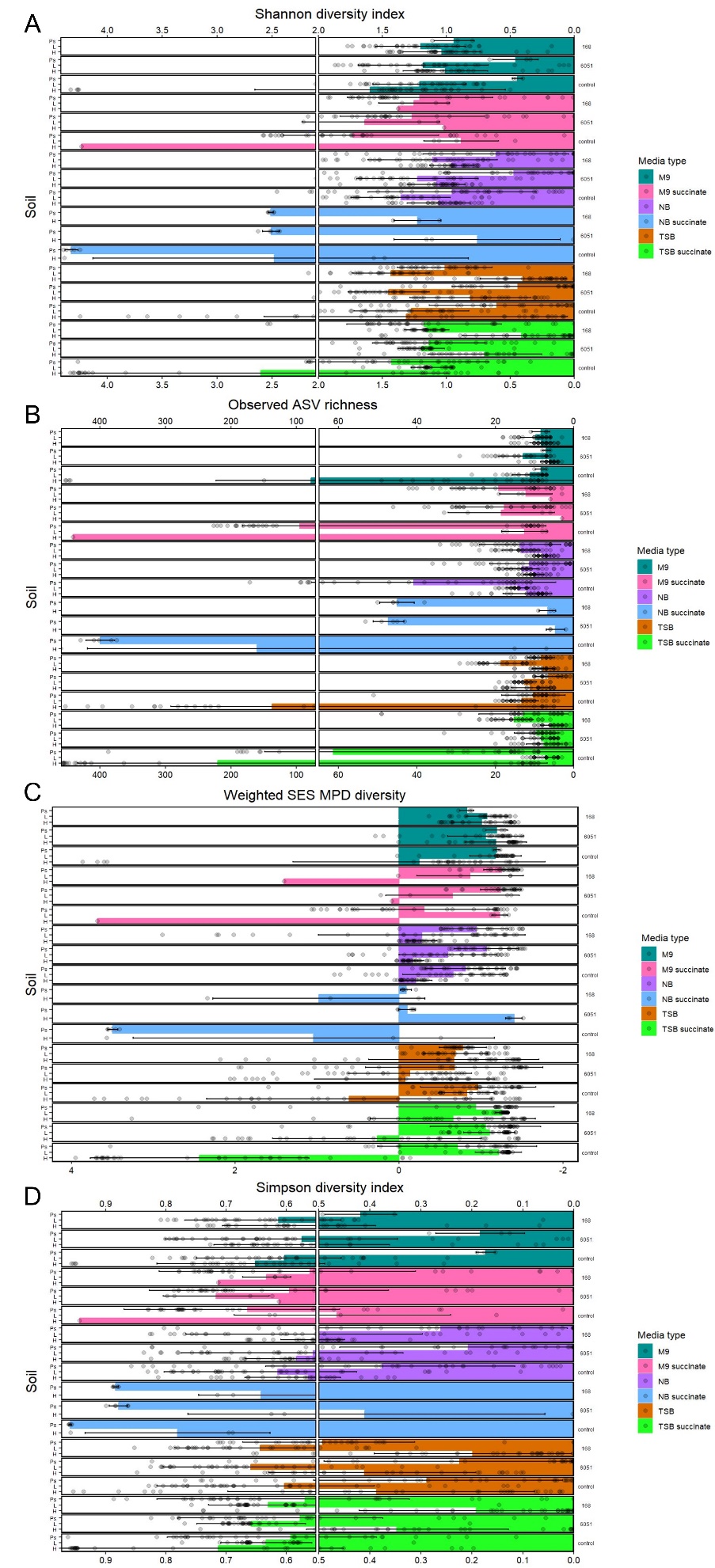

**Figure 19**. **Overview of impact on SynComs from inoculation process using alpha diversity and richness metrics**

After analyzing all the SynComs (control or base and inoculation conditions) post 16S rRNA sequencing, several metrics were calculated to assess the change in the SynComs after inoculating the SynComs with 168 and 6051a (6051) strains. Shannon diversity index (A), standardized effect size (SES) of mean pairwise distance (MPD) phylogenetic diversity (C) and Simpson diversity index (D) are shown for alpha diversity and Observed amplicon sequencing variant (ASV) richness (B) shown for estimating richness. Inoculation type is shown on the right side of the plots and soil type is shown on the left side with abbreviations denoted as H (Hopland), L (LLNL) and Ps (Potting soil). Data for alpha diversity is mean ± SD with each SynCom represented as a point which is up to 32 individual replicates as there were 16 SynComs in each media type of every soil and two separate 96-well plates were sequenced for every condition.

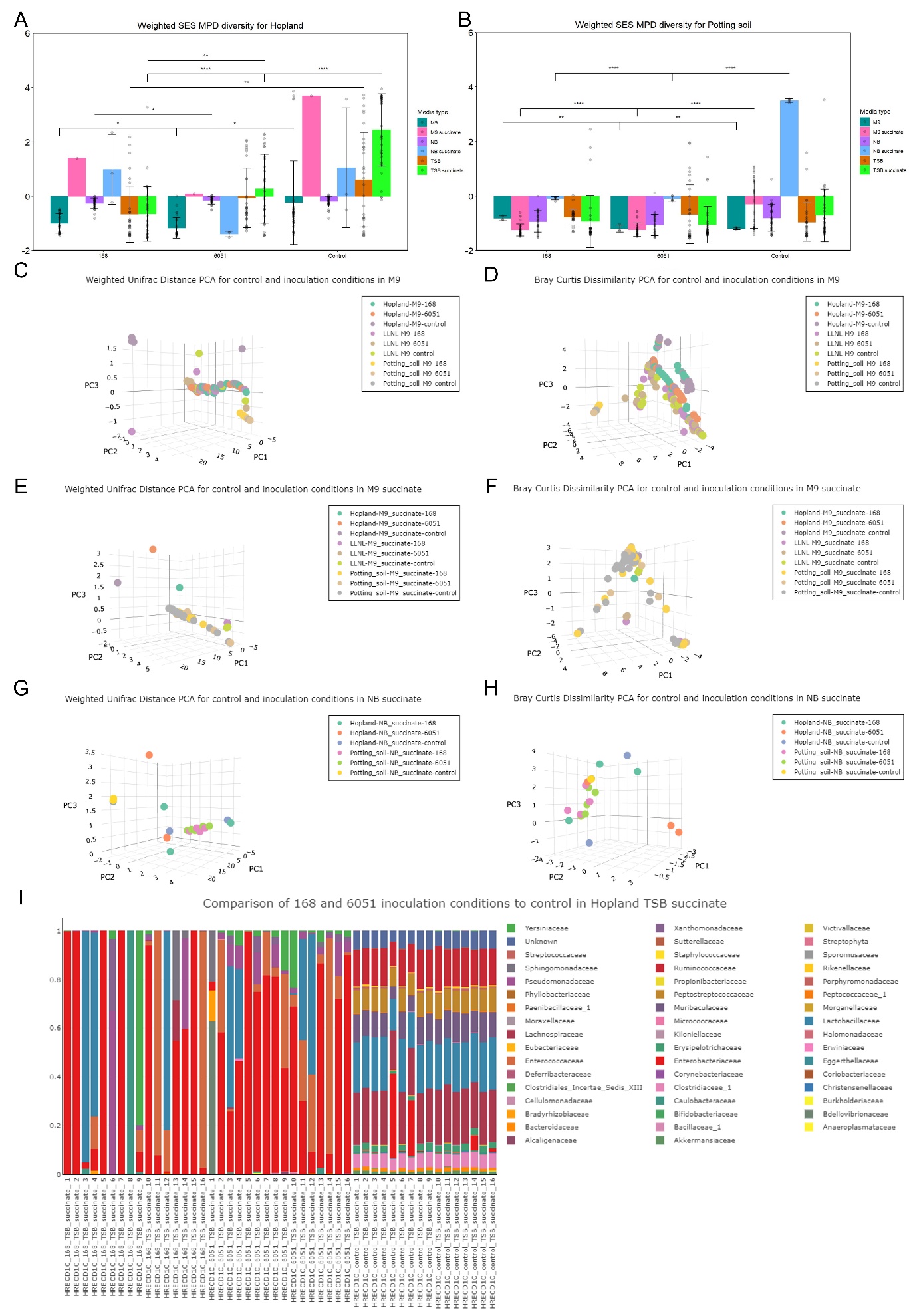

**Figure 20**. **Further assessment of how SynComs in Potting soil and Hopland were altered by attempting to incorporate *B. subtilis* 168 and 6051a strains**

After analyzing all the SynComs (control or base and inoculation conditions) post 16S rRNA sequencing, several metrics were calculated to assess the change in the SynComs after inoculating the SynComs with 168 and 6051a (6051) strains. Standardized effect size (SES) of mean pairwise distance (MPD) phylogenetic diversity is shown for alpha diversity (A, B) with inter-inoculation statistics shown. For beta diversity, Weighted UniFrac distance (C, E, G) and Bray Curtis Dissimilarity (D, F, H) are visualized as PCA plots with graphs grouped based on media type then comparing soil and inoculation type. A representative composition profile at the family level (I) is shown to exemplify the impact from the inoculation process. Data for alpha diversity is mean ± SD with each SynCom represented as a point which is up to 32 individual replicates as there were 16 SynComs in each media type of every soil and two separate 96-well plates were sequenced for every condition with the same for PCA plots; *p<0.05, **p<0.01, ****p<0.0001.

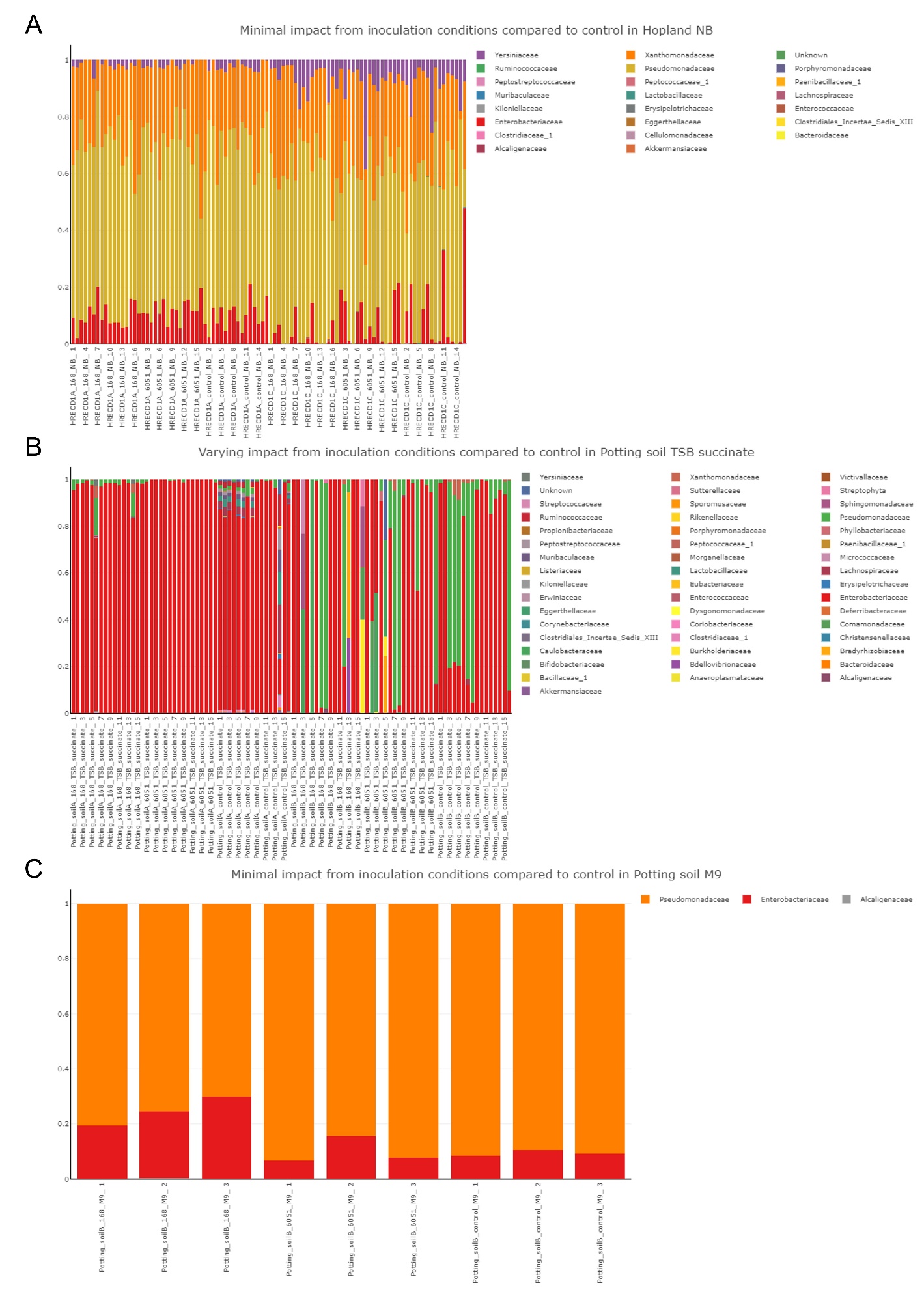

**Figure 21**. **Representative composition profiles highlight the lack of impact from the inoculation process on SynComs from different soil and media types**

Three representative composition profiles at the family level were assembled to display various trends observed in the SynComs and contextualize the results seen in the other metrics. The designation of control is applied to these SynComs as these are the base SynComs that the two *B. subtilis* strains were inoculated into generating the other SynComs. The first compositional profile was used to show lack of change in community structure in Hopland (HRECD1) NB after the 168 and 6051a (6051) strains were inoculated and compared to the base SynComs (A). The second compositional profile was used to represent SynComs that showed minimal change in community structure in Potting soil TSB succinate after the inoculation process (B). The third compositional profile was used to represent SynComs that showed minimal change in community structure in Potting soil M9 after the inoculation process (C).

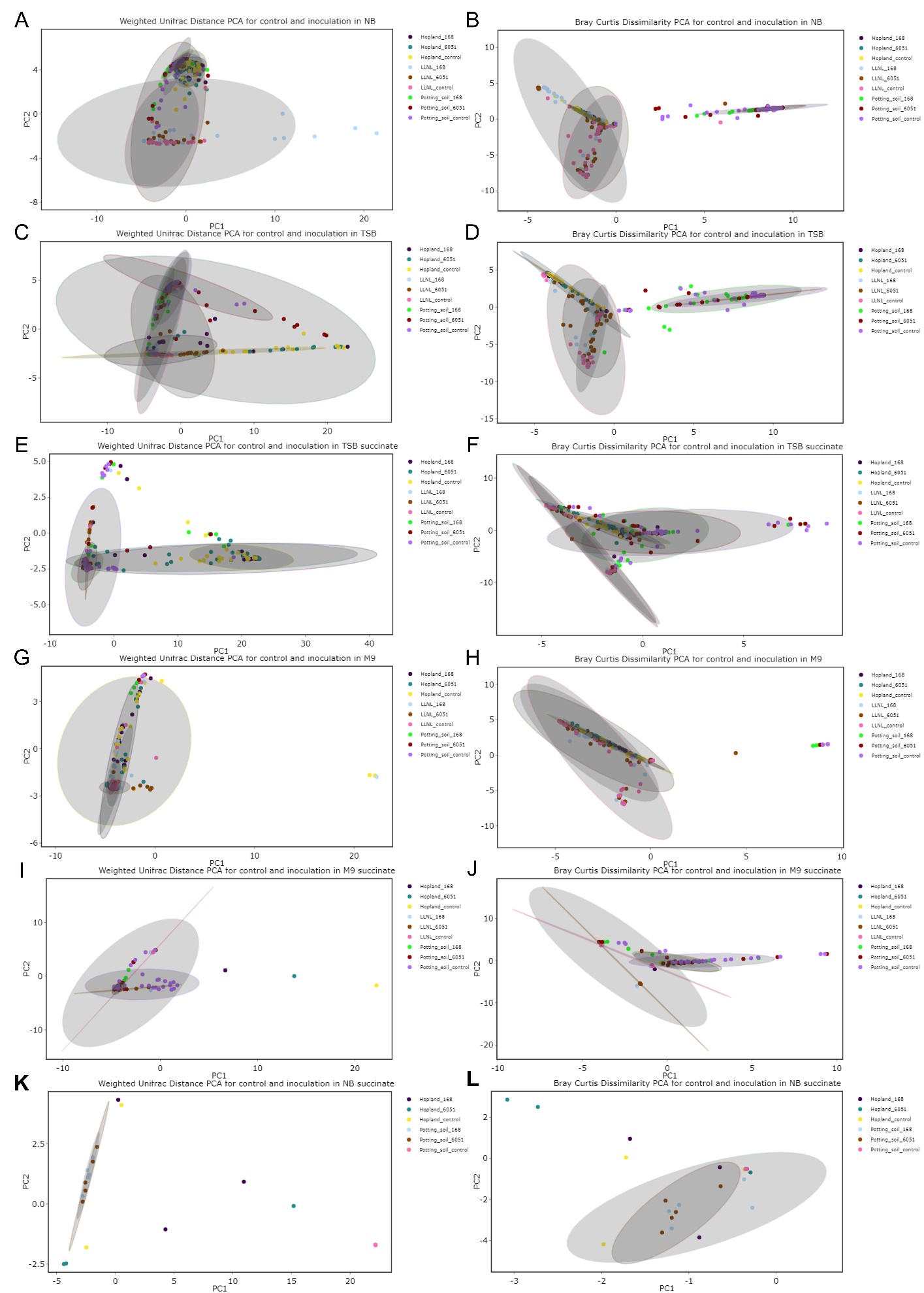

**Figure 22**. **Overview of impact on SynComs from inoculation process using beta diversity metrics**

Weighted UniFrac distance (A, C, E, G, I, K) and Bray Curtis Dissimilarity (B, D, F, H, J, L) are shown for beta diversity as PCA plots with the ellipses representing 95% confidence intervals where the plots are structured based on media type so that soil and inoculation conditions can be compared. Data is composed of up to 32 individual replicates as there were 16 SynComs in each media type of every soil and two separate 96-well plates were sequenced for every condition.

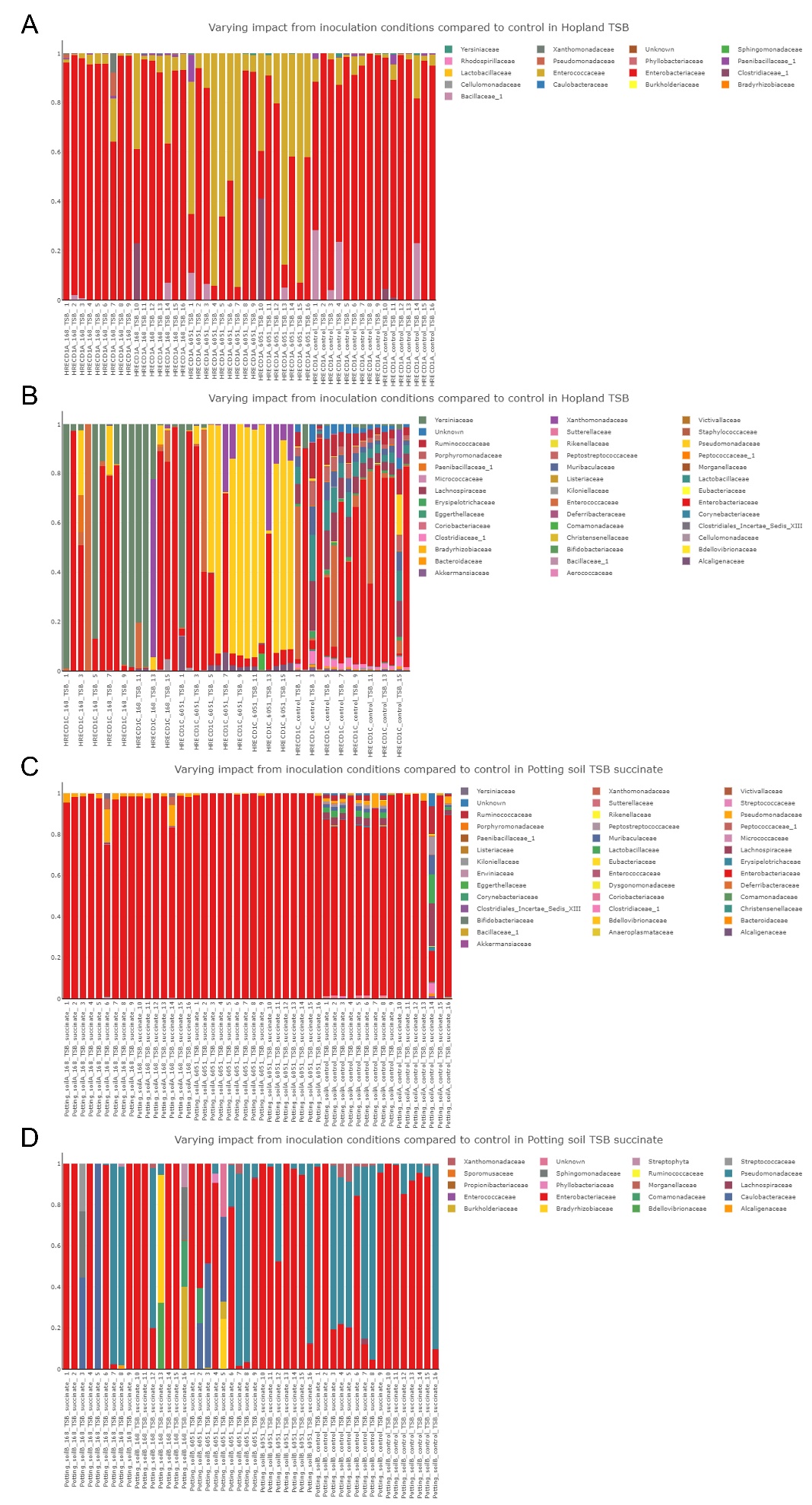

**Figure 23**. **Representative composition profiles highlight the variance of impact between well-plates from the inoculation process on SynComs from different soil and media types**

Four representative composition profiles at the family level were assembled to display various trends observed in the SynComs and contextualize the results seen in the other metrics. The designation of control is applied to these SynComs as these are the base SynComs that the two *B. subtilis* strains were inoculated into generating the other SynComs. The first set of compositional profiles were used to show variance of change in community structure between well-plates in Hopland (HRECD1) after the 168 and 6051a (6051) strains were inoculated and compared to the base SynComs (A, B). The second set of compositional profiles were used to show variance of change in community structure between well-plates in Potting soil after the 168 and 6051a (6051) strains were inoculated and compared to the base SynComs (C, D).

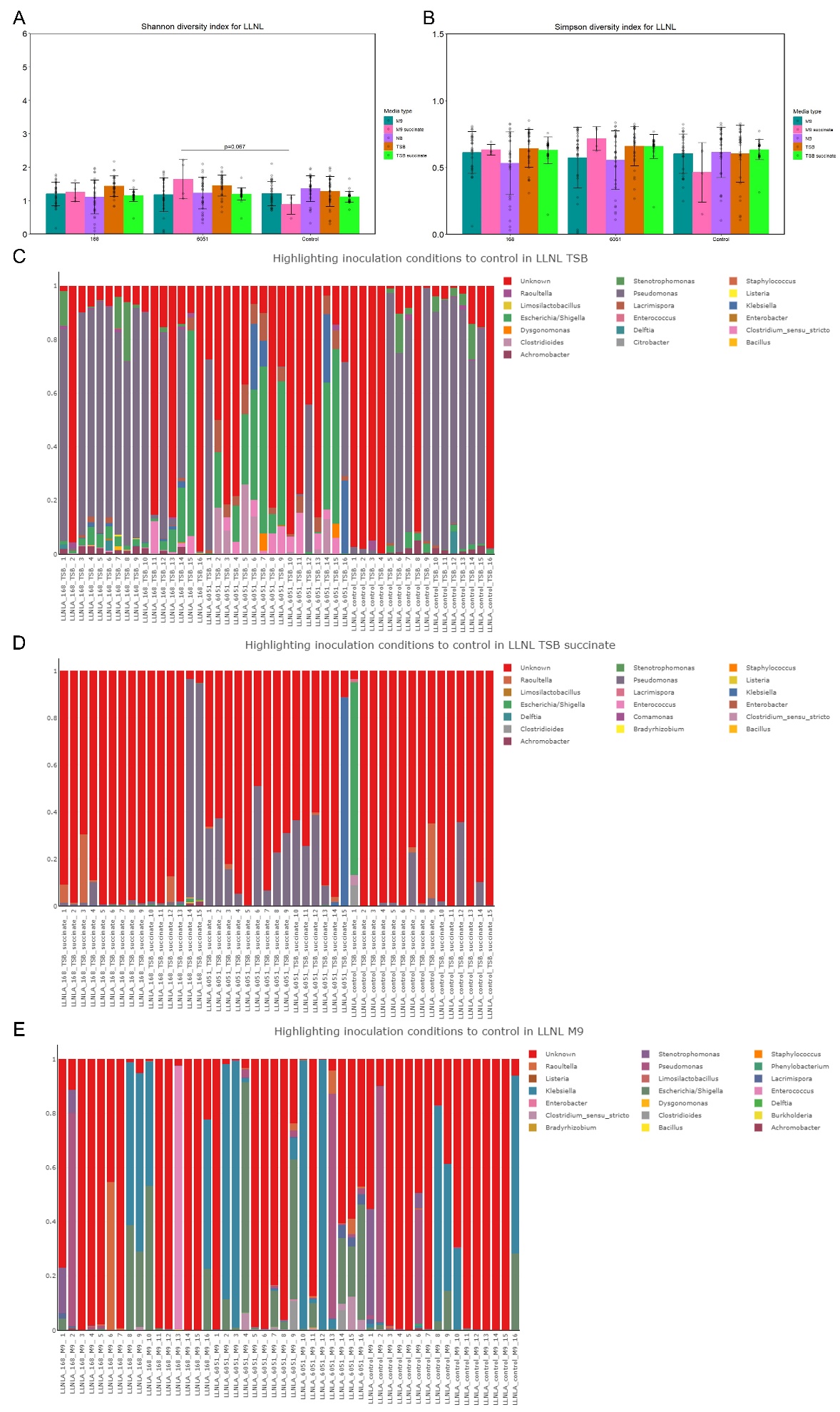

**Figure 24**. **Further assessment of impact from inoculation process compared to control SynComs in LLNL conditions**

After analyzing all the SynComs (control or base and inoculation conditions) post 16S rRNA sequencing, several metrics were calculated to assess the change in the SynComs after inoculating the SynComs with 168 and 6051a (6051) strains. Shannon diversity index (A) and Simpson diversity index (B) are shown for alpha diversity metrics. Three representative composition profiles at the genus level were assembled to display various trends observed in the SynComs and further contextualize the results seen in the other metrics during inoculation conditions. The compositional profiles were used to show community structure in LLNL for 168 and 6051a (6051) which included TSB (D), TSB succinate media type (D)and M9 media type (E). Data for alpha diversity is mean ± SD with each SynCom represented as a point which is up to 32 individual replicates as there were 16 SynComs in each media type of every soil and two separate 96-well plates were sequenced for every condition.

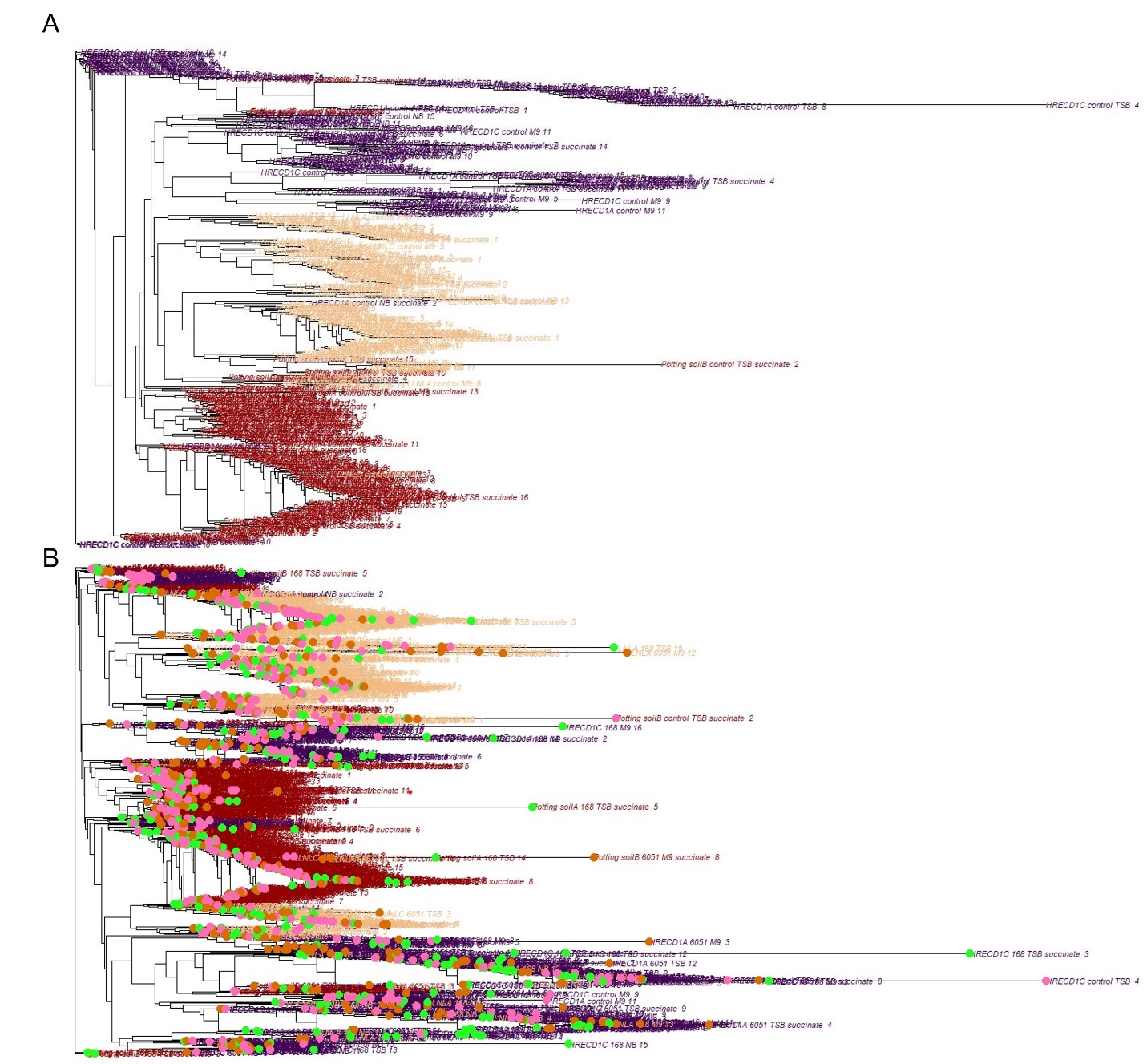

**Figure 25**. **Phylogenetic trees at the community level emphasize impact of the inoculation process on the structure and grouping of the SynComs**

The base or control top-down SynComs were constructed into rooted phylogenetic tree at the community level (A) to compare to a phylogenetic tree of all SynComs including the inoculated SynComs (B). Both phylogenetic trees were colored based on soil type with Hopland (HRECD1) colored in purple, LLNL colored in beige and Potting soil colored in maroon. The second tree (B) was constructed where the soil types are still colored the same as the first then the inoculation type was overlayed onto the tree using colored dots: control or base SynCom (pink), 168 strain (light green) and 6051a (6051) strain (orange).

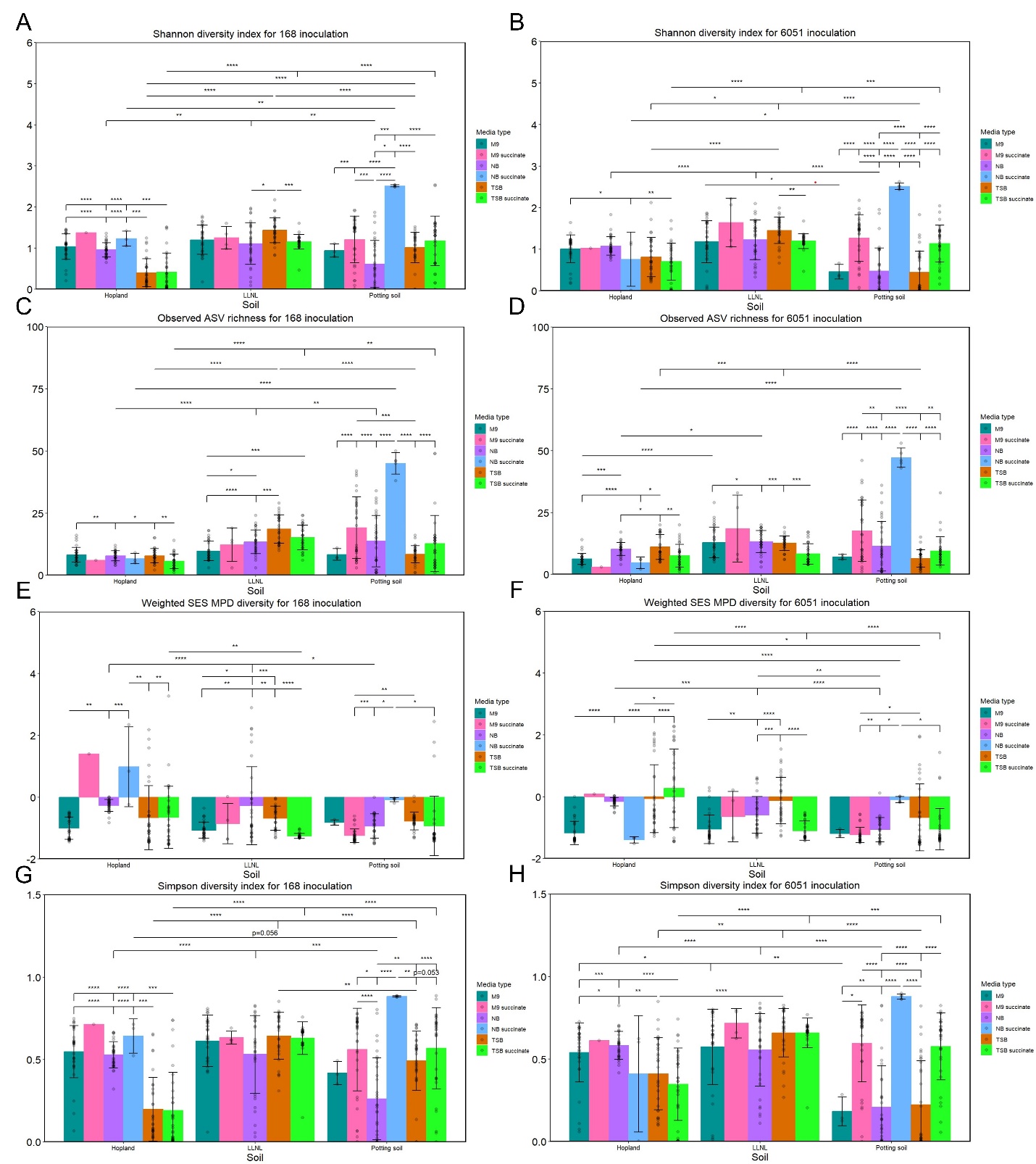

**Figure 26**. **Further assessment of how *B. subtilis* 168 and 6051a strains inoculation resulted in variable outcomes for complexity of SynComs based on soil and media type**

After analyzing all the inoculated SynComs post 16S rRNA sequencing, several metrics were calculated to assess the change in the SynComs after inoculating the SynComs with 168 and 6051a (6051) strains. Shannon diversity index (A, B), standardized effect size (SES) of mean pairwise distance (MPD) (E), SES mean nearest taxon distance (MNTD) phylogenetic diversity (F) and Simpson diversity index (G, H) are shown for alpha diversity with Observed amplicon sequencing variant (ASV) richness (C, D) used for estimation of richness with intra- and inter-soil statistics shown on the same graph. The 168 conditions are shown on the left (A, C, E, G) and 6051a conditions are on the right (B, D, F, H). Data for alpha diversity is mean ± SD with each SynCom represented as a point which is up to 32 individual replicates as there were 16 SynComs in each media type of every soil and two separate 96-well plates were sequenced for every condition; *p<0.05, **p<0.01, ***p<0.001, ****p<0.0001.
